# A simple immunoassay for extracellular vesicle liquid biopsy in microliters of non-processed plasma

**DOI:** 10.1101/2021.09.20.461033

**Authors:** Carmen Campos-Silva, Yaiza Cáceres-Martell, Estela Sánchez-Herrero, Amaia Sandúa Condado, Alexandra Beneitez-Martínez, Álvaro González Hernández, Mariano Provencio, Atocha Romero, Ricardo Jara, María Yáñez-Mó, Mar Valés-Gómez

**Author notes:** Corresponding author: Mar Valés-Gómez. Department of Immunology and Oncology, National Centre for Biotechnology, CNB-CSIC, Madrid, Spain.

## Abstract

Extracellular vesicles (EVs), released by most cell types, provide an excellent source of biomarkers in biological fluids. Here we describe a method that, using just a few microliters of patient’s plasma, identifies tumour markers exposed on EVs. Studying physico-chemical properties of EVs in solution, we demonstrate that they behave as stable colloidal suspensions and therefore, in immunocapture assays, many of them are unable to interact with a stationary functionalised surface. Using flocculation methods, like those used to destabilize colloids, we demonstrate that cationic polymers increase EV ζ-potential, diameter, and sedimentation coefficient and thus, allow a more efficient capture on antibody-coated surfaces by both ELISA and bead-assisted flow cytometry. These findings led to optimization of a protocol in microtiter plates allowing effective immunocapture of EVs, directly in plasma without previous ultracentrifugation or other EV enrichment. The method, easily adaptable to any laboratory, has been validated using plasma from lung cancer patients in which the epithelial cell marker EpCAM has been detected on EVs. This high throughput, easy to automate, technology allows screening of large numbers of patients to phenotype tumour markers in circulating EVs, breaking barriers for the validation of proposed EV biomarkers and the discovery of new ones.

## INTRODUCTION

During their normal life cycle, cells can release different types of vesicles originated from a variety of processes involving membrane invaginations and pinching-off events ^1^. Thus, in biological fluids, different types of extracellular vesicles (EVs) can be found, providing information of the different physio-pathological processes occurring in any individual and allowing trafficking of diverse subcellular components which can act as mediators in the exterior milieu ^2-4^. Exosomes are one type of such extracellular nanovesicles, generated from the endocytic pathway. Since vesicle cargo reflects cell composition, EVs are regarded as a potential useful tool displaying biomarkers and so, they have attracted recently great attention from scientists, clinicians and biotechnological companies ^5-9^.

Multiple candidate EV biomarkers have been suggested for different pathologies, however, for a routine translation into the clinics, high throughput comparative studies are still necessary to define the suitability of each particular biomarker in a given disease context. Such association studies, requiring the analysis of samples from large patient cohorts, are hampered by the current available methods which require either relatively large sample volumes or long protocols of nanovesicle pre-enrichment together with the use of sophisticated equipment or specialised personnel ^10-13^. The development of nanosensors makes possible to specifically detect EVs using small sample volumes from large numbers of patients ^14-16^, but these new technologies require purpose-designed devices assembled in specialised laboratories and, data derived from this type of study are still scarce. Thus, carrying out large screening projects studying EVs in a research or clinical setting would require the adaptation of widely used techniques to allow the identification of vesicles in small volumes and with minimal sample manipulation. An extra level of complication arises from the fact that any biological fluid contains EVs from many cellular origins and samples can be very heterogeneous ^17^. Therefore, marker selection is paramount for the characterization of the composition, number and size of the different vesicle subpopulations that can be released by any cell ^2, 18-21^.

To simplify the detection of EV proteins using a technique readily available in most clinical settings, we recently defined the critical parameters for improved flow cytometry detection of EVs after immunocapture ^22^. To further improve these assays, we explore here the hypothesis that EVs are stable in suspension with the physico-chemical properties of colloids, in which EVs would correspond to the disperse phase and the buffer to the solvent of a colloidal suspension ^23-25^. In a colloidal suspension the particles do not sediment as a consequence of gravity, instead they move erratically in such manner that the electrostatic repulsion is significantly larger than the thermal energy, and molecular attraction forces, such as van der Waals, do not prevail ^25^. This could limit the encounter with functionalised surfaces, such as antibodies on micro-beads, that, in contrast to EVs, would rapidly sediment to the bottom of the tube. If this hypothesis is true, destabilising the EVs suspension might increase the detection capacity in immunocapture experiments.

The stability of colloid suspensions has been studied in detail, in particular in several industrial settings such as wastewater treatment. Depletion mechanisms and other destabilising factors including the addition of external particles, such as polymers, cause precipitation of the colloidal suspension, an effect known as flocculation ^26, 27^.

Here we analysed the effects on EV suspensions of two cationic polymers: poly-L-lysine (PL) and hexadimethrine bromide (PB) (polybrene, commercial brand name) commonly used to increase the efficiency of drug and virus delivery to cells via aggregation, sedimentation and adsorption ^28^. We demonstrate that the addition of positively charged polymers to EV suspensions leads to the precipitation of vesicles that enhances antibody capture of EVs in both ELISA and bead-based assays.

Thus, by taking advantage of the colloidal properties of EVs, we have developed a method combining cationic polymers with the maximization of EV-surface contact to directly phenotype tumour antigens contained in nanovesicles from patient biological samples. Using immunocapture in bead-assisted flow cytometry, tumour markers were easily detected in only a few microliters of body fluids, without previous ultracentrifugation or enrichment of vesicles. These new experimental conditions, that radically improve the efficiency of EV detection in immunocapture assays, open new possibilities for the study of samples from large cohorts of patients and controls with minimal effort in any laboratory setting.

## MATERIALS AND METHODS

### Cells lines and reagents

Metastatic Melanoma cell lines: Ma-Mel-55, Ma-Mel-86c, (derived from melanoma patient metastases), provided by Prof. Annette Paschen (University Hospital of Essen, Germany), have been described elsewhere ^29-31^. Lung cancer cell line H3122 from ATCC was authenticated by satellite analysis at the genomics service of the Institute of Biomedical Research (IIB-CSIC). Cells were regularly assayed for mycoplasma contamination.

Metastatic Melanoma and lung cancer cell lines were cultured in RPMI 1640 (Sigma-Aldrich Co., St Louis, MO, USA) with 10% fetal bovine serum (FBS), 1 mM L-Glutamine, 1 mM Sodium Pyruvate, 0.1 mM non-essential amino acids, 10 mM HEPES and Penicillin and Streptomycin at 100 μg/ml, at 37°C, in 5% CO_2_/95% air, and passaged when cells reached 80-90% confluence.

Unless otherwise stated, all chemicals were purchased from Merck & Co (Kenilworth, New Jersey, USA), including hexadimethrine bromide ≥94% also known as polybrene, and poly-L-lysine solution – 0.1%(w/v) in H_2_O.

Antibodies used for Western Blot include mouse monoclonal anti β-actin (clone AC-15, Sigma, St. Louis, MO, United States) at 0.13 μg/ml; anti tetraspanins: anti CD81 (clone M-38 kind gift from Vaclav Horejsi, Croatia), anti CD9 (clone VJ1/20), anti CD63 (clone Tea3/18); biotinylated anti-EpCAM (clone VU-1D9) (all from Immunostep S.L, Salamanca, Spain); all used at 1 μg/ml; and biotinylated goat polyclonal anti-MICA antibody (BAF1300, R&D Systems, Minneapolis, Minnesota, United States) at 2 μg/ml. For capture in immunoassays, antibodies used were monoclonal mouse anti-MICA (MAB13002, R&D biosystems), anti-EpCAM (clone VU-1D9), anti-CD63 (clone Tea3/18) (Immunostep S.L. Salamanca, Spain) or IgG1 (MOPC 21, Sigma, St. Louis, MO, USA), as isotype control. For detection in immunoassays monoclonal mouse anti-CD81 (clone M38), anti-CD9 (Clone VJ1/20) (Immunostep, S.L., Salamanca Spain) and IgG1 (MOPC-21, Biolegend, San Diego, California, USA), all directly conjugated to PE were used at 0.02 μg/μl.

### EV-enriched preparations

Cells were grown until 70% confluence and then changed into medium prepared with 1% EV-free FBS (prepared by ultracentrifugation at 100,000 × *g* for 20 hours), for EV accumulation during 3-4 days. Cell supernatants were centrifuged for 10 min at 200 × g and small EVs enriched by sequential centrifugation as previously described ^32, 33^. After ultracentrifugation at 100,000 × *g* for 2 hours at 4°C, EVs were resuspended in 0.22 µm filtered HEPES-buffered saline (HBS: 10mM HEPES pH 7.2, 150mM NaCl) (2.67 µl/ml of starting cell culture supernatant) and stored at −20°C, for short term use, or at −80°C. For longer storage, EVs were lyophilised using a VirTis Freezemobile 12SL Freeze Dryer Lyophilizer (VirTis, Habour Group, St. Louis, MO, USA). Note that this protocol is used for exosomes enrichment, however, their specific origin cannot be assured so we refer to the enriched particles as EVs. For plasma and ascitic liquid EV enrichment, 200 µl of sample were diluted in 4 ml of HBS and ultracentrifuged at 110,000 × g for 2 hours at 4°C, the EV pellet was resuspended in 15 µl of HBS and stored at −20 °C for short term use.

### EV quantitation

The concentration and size of enriched EVs in each preparation were determined by nanoparticle tracking analysis (NTA) in a Nanosight NS500 (Malvern Instruments Ltd, Malvern, UK) equipped with a 405 nm laser, a sCMOS camera and NTA 3.1 Software. Enriched EV preparations were diluted (1:1000) for measurement at a concentration range of 1 × 10^9^ particles/ml, as recommended by manufacturer. The settings used were: Camera level:12, Threshold: 10, Capture: 60 s., Number of Captures: 3, Temperature 25 °C. The experiments were carried out at the laboratory of Dr. H Peinado, Spanish National Centre for Oncological Research (CNIO). Some measurements were double checked in a Zetaview (Particle Metrix).

### Electron Microscopy

For Transmission Electron Microscopy (TEM), 1 µl of the EV preparation obtained after sequential centrifugation was diluted 1:10 in filtered (0.22 µm) HBS. Ma-Mel-55 melanoma-derived EVs (0.6 × 10^9^ particles /µl), were used for the experiment with polymers. Polymers were added at a final concentration of 4 µg/ml of polybrene or Poly-L-lysine and incubated for 18 hours. Samples were floated on carbon-coated 400-mesh 240 Formvar grids, incubated with 2% uranyl acetate, and analysed using a Jeol JEM 1011 electron microscope (JEOL, Akishima, Tokyo, Japan) operating at 245 100 kV with a CCD camera Gatan Erlangshen ES1000W. The experiment was carried out at the Electron Microscopy Facility, Spanish National Centre for Biotechnology (CNB).

### Western and dot blots

EV enriched preparations (either 6.8 × 10^9^ particles or 5 μl of patient plasma EV preparation; 5.86 × 10^8^ H3122 EVs were used as EpCAM positive control) or respective cell lysates (30 μg) were loaded in 12% SDS-PAGE gels, either under reducing or non-reducing conditions, as indicated in the experiments, and transferred to membranes with Trans-Blot® Turbo™ Transfer Packs (Biorad, Hercules, California, USA). Membranes were blocked using 5% non-fat dry milk in PBS containing 0.1% Tween-20 (PBS-T). Primary antibody was incubated for 1 h in PBS-T and, after washing, membranes were incubated with the appropriated secondary antibody. Secondary antibodies used were Alexa-700 GAM or Alexa-790-SA (ThermoFisher), when proteins were visualised using the Odyssey Infrared system (LI-COR Biosciences, Lincoln, NE, USA). When proteins were visualised using the ECL system (Amersham Biosciences, Amersham, UK), horseradish peroxidase-conjugated goat anti-mouse antibody (Sigma, St. Louis, MO, USA) was used at 0.8 μg/ml or horseradish peroxidase-streptavidin (Biolegend, San Diego, California, USA) at 0.1 μg/ml. For dot blots, 1 μl of EVs were immobilised onto nitrocellulose membranes, blocked and developed as for WB membranes.

### Antibody coated magnetic beads preparation

Antibody-coated magnetic beads were obtained from the Exostep™ kit (Immunostep, S.L., Salamanca, Spain) or prepared by amine coupling capture antibodies onto magnetic fluorescent beads (APC and PerCP) either from Luminex (MagPlex Microsphere) or Bangs (QuantumPlex M SP Carboxil) as previously described (Campos-Silva *et al*., 2019, Scientific Reports 9:2042, 1-12).

### EV detection by Flow cytometry

EV samples were incubated with 3000 antibody-coated beads in either 100 μL, 50 μL or 12 μL of PBS containing 1% casein (EV enriched preparations diluted 1:100-1:1000) for 18 h, in either a 5 ml tube or a well in a 96 flat-bottom microtiter plate, as specified in each experiment, without agitation at room temperature (RT). Unless otherwise stated, when using biological samples, 12 μL of precleared samples were added to 12 μL of beads in PBS-casein (PBS containing 1% casein, Bio-rad Laboratories, Hercules, California, USA). Background signals were determined by comparison with antibody isotype-coupled beads or PE-conjugated isotype antibody, as specified in each experiment. Cationic polymers were added and mixed before the 18-hour incubation, unless otherwise stated, at a final concentration of 4 µg/ml of polybrene or poly-L-lysine. Control samples were incubated in the same volume of PBS-casein without polymer. After the capture step, beads were washed with PBS-casein and recovered using a Magnetic Rack (Ref Z5343, MagneSphere(R) (Promega, Madison, Wisconsin, USA), for tubes, or 40-285 Handheld Magnetic Separation Block (Millipore, Burlington, MA, USA), for 96 well plates. The recovered beads were stained with PE-conjugated anti-tetraspanin detection antibodies (at 0.02 µg/µl) during 1 hour at 4 °C. After antibody binding, beads were washed with filtered PBS, and recovered using the Magnetic Rack. Beads were acquired by flow cytometry using Gallios, Cytomics FC 500 (Beckman Coulter) or CytoFLEX (Beckman Coulter) and data were analysed using Kaluza (Beckman Coulter) or FlowJo (Tree Star, Inc) software. Single beads were gated in Forward Scatter in the region corresponding to 6 μm [established using calibration beads (FlowCheck ProTM fluorospheres, Beckman Coulter, Brea, CA, USA)], excluding bead doublets and selecting APC-positive events ^33^. PE MFI was analysed within the 6 μm-APC-positive events.

### Dynamic Light Scattering (DLS)

For ζ-potential measurement, EVs enriched from Ma-Mel-86c melanoma cells (1.75 × 10^9^ particles/µl), were diluted to a final concentration of 3.5 × 10^10^ EVs/ml in HBS (in the concentration range recommended for measurement by the instrument software) and treated, for 5 min or 18 h, with 8 µg/ml of cationic polymer polybrene or 4 µg/ml Poly-L-lysine (a non-treated control sample was prepared in parallel as a control), for 5 min or 18h. The samples were then loaded in Zetasizer Nano DTS 1070 cuvettes for ζ-potential measurement at NanoZS (Red badge) ZEN3600 (Malvern Pranalytical, Malvern, UK) at 25 °C.

For size distribution measurement, EVs enriched from Ma-Mel-86c melanoma cells (1.9 × 10^9^ particles/µl), were diluted to a final concentration of 1.9 × 10^9^ EVs/ml in HBS (in the range recommended for measurement by the instrument software). EVs were treated, for 5 min or 18 h, with 8 µg/ml of cationic polymer polybrene or 4 µg/ml Poly-L-lysine (a non-treated control sample was prepared in parallel as a control) and loaded in a ZEN0040 disposable cell for diameter measurement using the same Zetasizer NanoZS (Red badge) ZEN3600 equipped with a 633 nm laser. Readings were performed at 25°C.

DLS experiments were carried out at the Instituto de Ciencia de Materiales de Madrid - ICMM – CSIC. Malvern Pranalytical DTS Software Version 5.10 was used for data processing and analysis.

### Analytical ultracentrifugation and sedimentation coefficient analysis

For analytical centrifugation experiments, Ma-Mel-86c-derived EVs (2.6 × 10^9^ particles/µl) were used, either directly after ultracentrifugation (diluted 1:10 in HBS) or subjected to further purification by size exclusion chromatography (SEC) to rule out any effect of co-precipitating proteins. For SEC, 26 µl of the EVs resuspended in HBS (pH 7.2, 0.22 µm filtered) after sequential centrifugation were layered on a 1 ml bed of sepharose CL-2B (Sigma CL2B300) in the same buffer. The eluate was collected in 0.2 ml fractions. Protein concentration was determined in each fraction, by measuring absorbance at 280 nm and the EV-containing fraction was determined by dot blot using anti-CD81 antibody. SEC was repeated 3 times and the EV-enriched fractions were pooled and used for sedimentation rate analysis experiments.

Both types of samples were treated either with 4 µg/ml of polybrene or 4 µg/ml of Poly-L-lysine (a non-treated sample was prepared in parallel as a control). After an 18-hour incubation, all samples were run in an XLI analytical ultracentrifuge (Beckman Coulter, Brea, CA, USA) (wavelength: 280, 655 × g (3000 rpm), 80 minutes, 20 °C). The results were analysed with SEDFIT 16.1c analysis Software with a confidence interval variation of the analysis of 0.68. The experiment was carried out at the laboratory of analytical ultracentrifugation and macromolecular interactions (LUAIM) at Centro de Investigaciones Biológicas Margarita Salas (CIB-CSIC).

### ELISA

Plates were coated with capturing antibodies at 6 µg/ml in BBS (Borate Buffered saline) overnight at 4°C. After blocking the plates with 1% casein-PBS for 2 h at 37°C, samples were added: EV enriched preparation diluted 1:100-1:1000 in PBS-casein and incubated 18 hours at RT. When adding cationic polymers, they were mixed with the sample, at a final concentration indicated in each experiment, before the 18-hour incubation. PBS-casein was added to controls to maintain the same final volume. Biotinylated secondary anti-tetraspanin antibodies were added at 0.4 µg/ml and followed by 0.25 µg/ml streptavidin-HRP (Amersham). The reaction was developed using TMB (3,3⍰,5,5⍰-Tetramethylbenzidine) substrate (1-Step™ Ultra TMB-ELISA Substrate Solution; Thermo Scientific, Waltham, MA, USA). Absorbance was measured at 450 nm with Multiskan™ FC Filterbased Microplate Photometer (Thermo Scientific, Waltham, MA, USA).

### Healthy donor plasma and patient selection

Experiments were carried out following the ethical principles established in the 1964 Declaration of Helsinki. Patients (or their representatives) were informed about the study and gave a written informed consent. This study used samples from 2 hospitals in Spain, Clínica Universidad de Navarra and Hospital Universitario Puerta de Hierro. Samples from Hospital Universitario Puerta de Hierro were obtained through the development of the research projects “PI17/01977” and “PIE14/00064”. Both projects were approved by the Hospital Puerta de Hierro Ethics Committee (internal code 79-18 and PI144, respectively). Collection of cancer patient samples at the Clínica Universidad de Navarra was approved by the Clínica Universidad de Navarra Ethics Committee 111/2010 “Estudio traslacional prospectivo de determinación de factores predictivos de eficacia y toxicidad en pacientes con cáncer” and included in the Spanish National Biobank Register with the code C.0003132 (Registro Nacional de Biobancos). A small cohort including 24 lung cancer patients and 12 healthy donors was approved by the project 2021.145. Demographic and clinical data from lung cancer patients are available in Supplementary Table 1. Plasma, ascites and/or urine were collected from breast, ovarian and prostate cancer patients to test the methodology using different biological fluids.

Blood was collected from each subject in a 5 ml EDTA tube containing a gel barrier (PPT™, BECTON DICKINSON) to separate the plasma from blood cells after centrifugation. Plasma samples were frozen at −80°C until test. Ascites were centrifuged 10 minutes at 1500 × *g* after collection and frozen at −80°C until test. After the first thawing, aliquots were prepared and frozen to avoid further freezing-thawing cycles.

### Statistics

Graphpad Prism 8 software was used for statistical analysis and representation of the data. Statistical tests used are indicated in each figure legend.

## RESULTS

### Standard antibody-binding conditions are not 100% efficient for the capture of extracellular vesicles

We have recently described a high sensitivity method for immunocapture and detection of EVs by flow cytometry, based on the use of antibody-coated beads followed by detection with a labelled antibody ^22, Campos-Silva et al, Melanoma. Methods and protocols. Methods in Molec Biol. 2021^. During the optimisation of that method, we calculated the theoretical number of EVs that would bind the antibody-coated microspheres and analysed the saturation curve in experiments with increasing amounts of EVs. Although 6000 beads could theoretically bind 3.85 × 10^7^ EVs, our experimental data showed that saturation of detection occurred when around 3.6 × 10^9^ EVs (NTA measurement) were offered, suggesting that the beads were not capturing all the EVs that were present in the incubation mix. NTA measurement may overestimate the number of EVs, since the instrument does not discriminate protein aggregates from EVs, but a 2-log error in quantitation seemed improbable. Alternatively, since EVs are usually a heterogeneous mix, it could be possible that not all the EVs in the mixture contained the epitope for the capture antibody. To directly test these possibilities, nested rounds of incubations were carried out. EVs derived from a melanoma cell line were characterised (Figure 1 A, B, C) following MISEV2018 guidelines ^34^ and incubated with anti-CD63-coated beads for flow cytometry analysis. The supernatant from the first incubation was recovered and used with fresh anti-CD63-coated beads for successive flow cytometry analysis, until signal was minimal. Melanoma EVs were still detected on supernatants after several rounds of capture using anti-CD63-coated beads (Figure 1 D), suggesting that not all the EVs carrying the epitope were captured in a single step. We could observe the same behaviour with EV samples from different cell lines (not shown).

**Figure 1.**
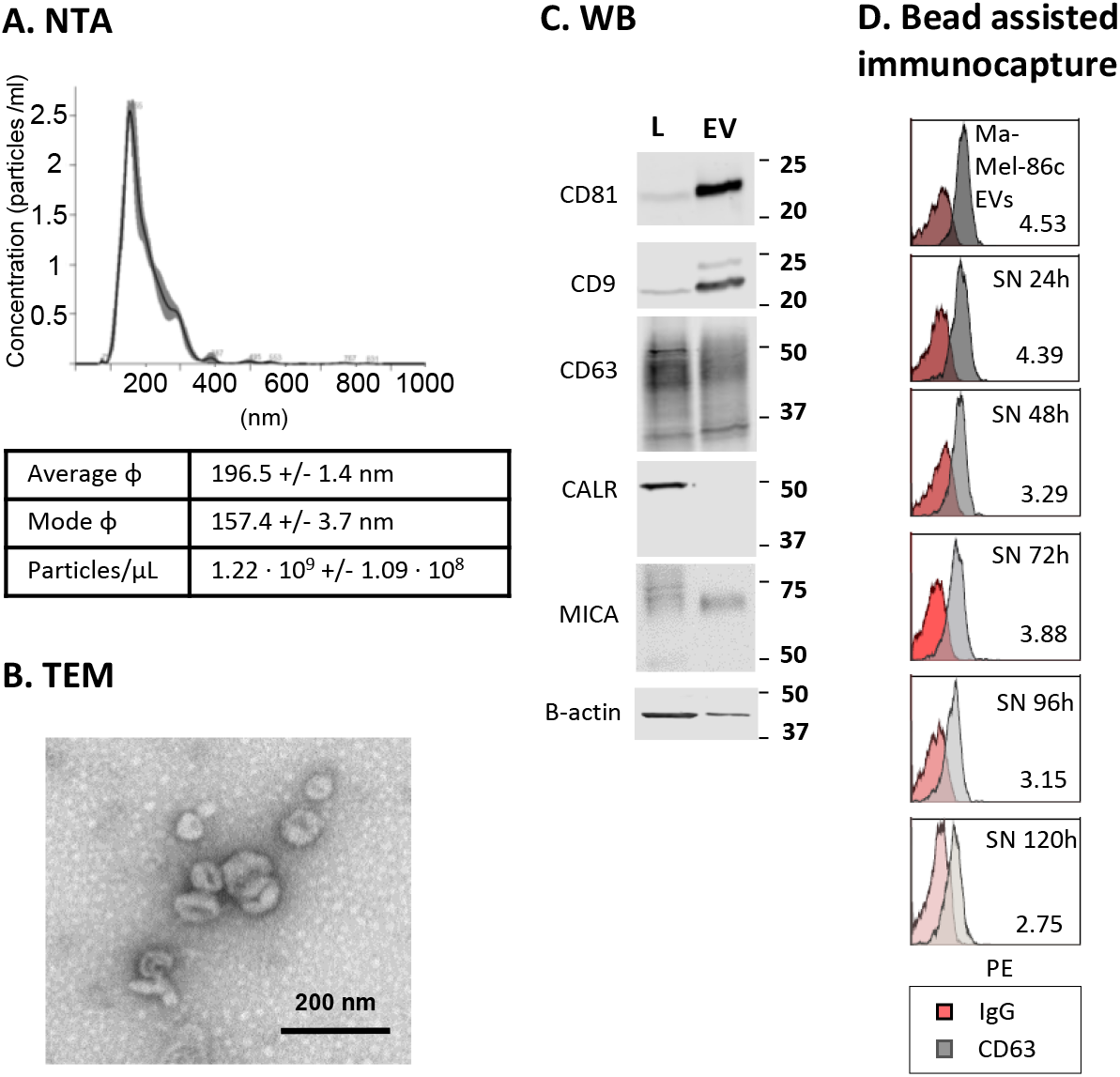
Repeated incubations are needed to immuno-capture all available EVs. Melanoma-derived EVs (from Ma-Mel-86c cell line) were enriched by ultracentrifugation. **A. Nanoparticle tracking analysis (NTA) of melanoma-derived EVs**. Average size and concentration of EVs were obtained in a Nanosight equipment capturing 3 videos of 60 s per measurement, with camera level 12, threshold 10 and temperature of 25 °C. Software NTA 3.1 (Malvern) was used for the analysis. ϕ: diameter. **B. Transmission Electron Microscopy (TEM)**. 1 µL EVs were diluted 1:10 in HBS and floated on a carbon-coated 400-mesh 240 Formvar grid, then incubated with 2% uranyl acetate and analysed using a Jeol JEM 1011 electron microscope operating at 245 100 kV with a CCD camera Gatan Erlangshen ES1000W. Pictures were taken at the Electron Microscopy Facility of the CNB. Bar: 200 nm. A representative image is shown. **C. EV characterization by Western Blot**. Whole cell lysates (L) and EVs were loaded in 12% SDS-PAGE gels. Membranes were immunoblotted for detection of: tetraspanins CD9, CD63, CD81 as general EV markers; β-actin as loading control; MICA as cancer-related marker; and calreticulin (CALR) as an endoplasmic reticulum resident protein not present in the EV fraction. Two gels were loaded: one gel, under non-reducing conditions and the other under reducing conditions, for actin detection. One representative experiment out of 3 is shown. **D. EV immuno-capture followed by flow cytometry**. 3000 anti-CD63-coated beads [or Isotype (IgG) coated as a negative control] were incubated with 3.5 × 10^6^ Ma-Mel-86c derived EVs/μl in 100 μl (3.5 × 10^8^ EVs/tube). Vesicles were detected by flow cytometry after incubation with anti-CD81-PE. Supernatants from the first incubation (SN 24h), containing unbound vesicles, were incubated again with fresh anti-CD63 beads and EVs captured during this second incubation were analysed by flow cytometry. This procedure was repeated the following 4 days (SN 48h, SN 72h, etc, as indicated). The histograms with the Relative Fluorescence Intensity 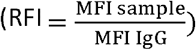 values from a representative experiment out of 3 is shown.

### EVs in solution form stable colloidal suspensions, which are destabilised using charged polymers

Agitation did not improve the efficiency of EV binding to beads, while a long incubation time was crucial for good detection ^22^. Thus, we decided to explore whether a more efficient capture of EVs could be achieved by affecting the physico-chemical properties of EVs. Because of their nanometric size, EV preparations could be considered colloidal suspensions, where EVs are stabilised in solution by steric (they are covered by proteins that would act as a solvated layer or halo) and electrostatic (amino acid and lipid charges) factors. Indeed, different methods initially developed for colloids, such as NTA, are used to characterise EV preparations ^35, 36^. Thus, experiments to identify the colloidal behaviour of EVs were carried out including their stability versus flocculation properties.

In order to evaluate the effect of depletion forces, several biophysical parameters were analysed after incubation of melanoma-derived EV suspensions with two cationic polymers, commonly employed in biology for precipitation of nanometric structures ^28^: hexadimethrine bromide (polybrene) and poly-L-lysine. First, the diameter and ζ-potential of metastatic melanoma derived EVs obtained by ultracentrifugation were measured by Dynamic Light Scattering (DLS) (NanoZS) (Figure 2A). Melanoma-derived EVs usually have a negative ζ-potential and this parameter can be used as an approximation to evaluate colloidal stability, since electrostatic repulsion prevents aggregation ^37^. When resuspended in regular isosmotic buffer, melanoma-derived EVs had, on average, a diameter of 196.5 nm by NTA. DLS readings of ζ-potential were −15.27 mV in average, while diameter measurements render a higher value of 302.97 nm with this technique. Interestingly, when EVs were incubated with 4-8 µg/ml of either polybrene or poly-L-lysine for 5 min, the ζ-potential of EVs increased to a range between −9.55 and +2.65 mV. Similarly, the average diameter of the EV suspension increased dramatically in the presence of the charged polymers when measured by DLS, accompanied by a higher poly-dispersity index (not shown). Longer incubation times resulted in bigger ζ-potential or diameter changes only in a few cases (polybrene at 4 µg/ml and poly-L-lysine at 4 µg/ml respectively). The observed trend was also confirmed by NTA (Zetaview technology, not shown).

**Figure 2:**
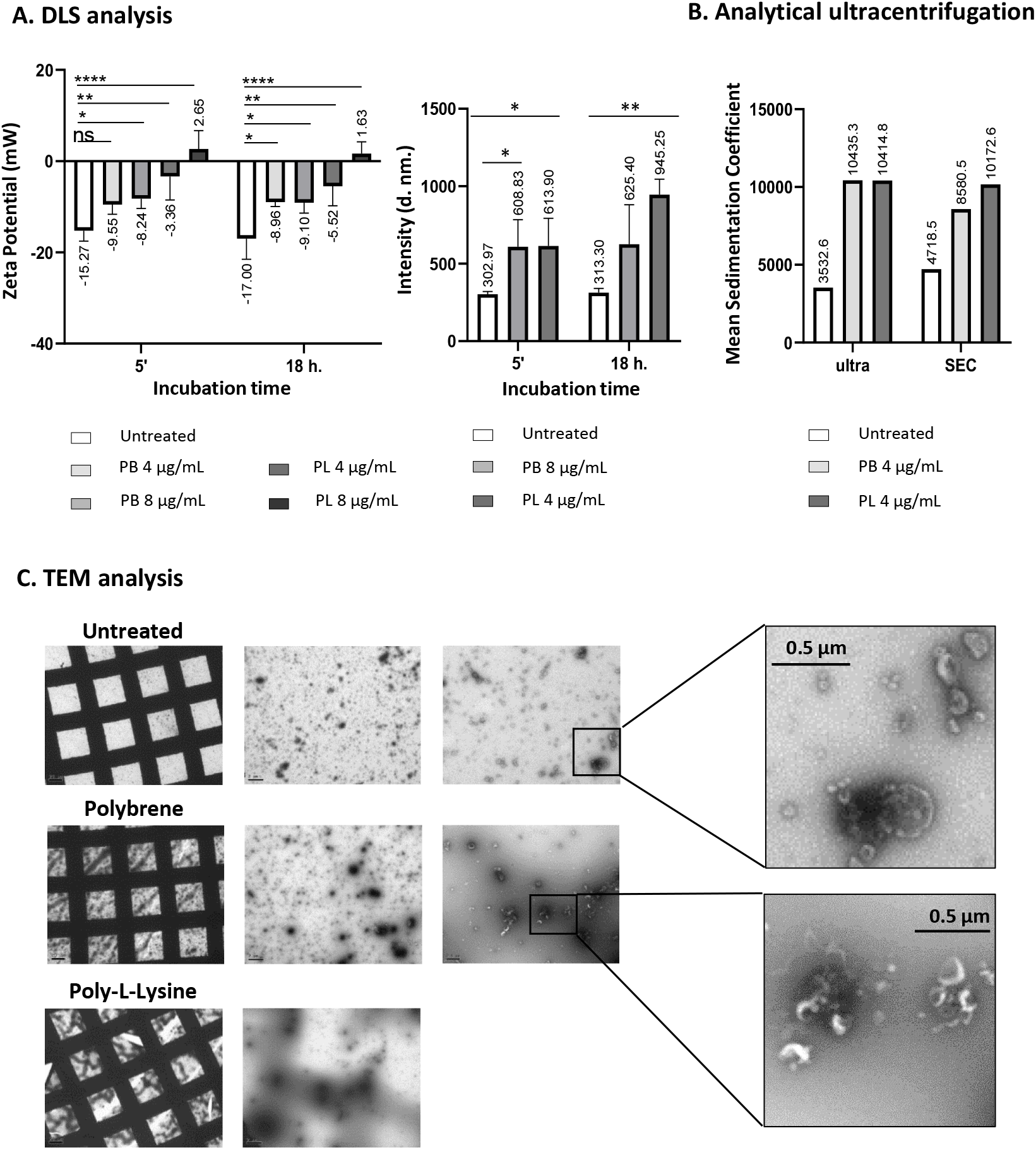
Cationic polymers affect the biophysical properties of EV suspensions and result in precipitation of the nanoparticles. Metastatic melanoma derived EVs obtained by ultracentrifugation were incubated with or without 4 μg/ml or 8 μg/ml Polybrene (PB) or 4 μg/ml or 8 μg/ml Poly-L-lysine (PL). **A. Zeta Potential and Hydrodynamic diameter by Dynamic Ligh Scattering (DLS)**. EVs were incubated either 5 minutes or 18 h with the polymers and analysed using a DLS instrument. Data on Zeta Potential and Intensity Mean (average diameter in nm.) are depicted. Mean and Standard Deviation from 3 independent experiments is shown. Statistical analysis was performed by a one-way ANOVA Fisher’s LSD test (p < 0.05). * p < 0.05, ** p < 0.01, *** p < 0.001, **** p < 0.0001). **B. Analytical ultracentrifugation**. The sedimentation coefficient of EVs was analysed either directly after ultracentrifugation (ultra) or after further purification by size exclusion chromatography (SEC). The graph represents data on the average sedimentation coefficient obtained after a 18 h-incubation with polymers of EVs obtained under these two methods. **C. Transmission Electron Microscopy**. The left column shows the general aspect of the grid, followed by sequential magnifications [Bar: 2 µm (2nd column), 0.5 µm (3rd and 4th columns)]. Polymers caused precipitation of EV samples, affecting the integrity of the resin upon electron beam incidence. Thus, pictures from Poly-L-Lysine treated samples could not be obtained at high magnification. Electron dense areas correspond to EV aggregates.

Since the diameter and charge data obtained by light scattering methods suggested that polymers could cause aggregation and flocculation of EVs, analytical ultracentrifugation was used to measure the sedimentation rate of nanoparticles in the presence of those positively charged polymers (Figure 2B). Indeed, the average sedimentation coefficient of the EV suspension obtained by ultracentrifugation increased dramatically in the presence of the charged polymers. To eliminate any possible contribution of proteins co-precipitating with EVs in ultracentrifugation, samples were further purified by SEC, yielding a similar result (Figure 2B). Interestingly, after an 18-hour incubation with cationic polymers, samples presented high polydispersity with multiple peaks displaying a wide range of sedimentation coefficients (not shown). This implies that larger particles of different sizes are present in the sample incubated with polymers compared to control samples. In line with these observations, electron microscopy imaging of EVs after an 18-hour incubation with cationic polymers revealed large high electron dense structures (1-2 µm), consistent with high mass particles (Figure 2C). In more detailed images, clusters of EVs could be observed. Inspection at low magnification of the grids already substantiated the high density masses. In fact, when the electron beam stroke on high density spots observed in poly-L-lysine-incubated samples, the resin ruptured and high magnification images could not be obtained.

Altogether, these results indicated that adding cationic polymers to EV preparations, generated aggregates of particles of different sizes, leading to higher sedimentation rates, confirming that EV suspensions behave as colloids.

### Cationic polymers increase the detection of EV proteins in immunoassays

Since cationic polymers destabilised EV suspensions, increasing the average diameter of the particles and the sedimentation coefficient, we hypothesised that this depletion force phenomenon, described for colloidal systems, could improve EV detection by immuno-capture methods. Thus, we tested how the inclusion of cationic polymers affected the precipitation of nanoparticles on antibody-coated surfaces either in bead-assisted flow cytometry or ELISA experiments. For these experiments, EVs obtained by ultracentrifugation from either the melanoma cell line Ma-Mel-86c or the lung cancer cell line H3122 were used. The characterization of these vesicles was performed first by classical methods including NTA, Western Blot and TEM to establish the tumour markers present on EVs derived from the different tumour cell lines (Figure 1 and Supplementary Figure 1). Ma-Mel-86c is positive for the tumour-associated immune ligand MICA as reported previously ^31^, while H3122 expresses the epithelial cell marker EpCAM (Supplementary Figure 1). Enhanced detection of both tetraspanins and tissue-specific markers was observed using 4 μg/ml of either polybrene or poly-L-lysine, as shown in the RFI values (Figure 3A), except for saturated signals (CD63-CD81 in Ma-Mel-86c EVs). Initial polymer titration experiments revealed that, in general, the best signal was obtained when polybrene was added at 8 µg/ml reaching a significant difference in the signal with respect to the untreated assay (Figure 3B). The same effect was observed in ELISA experiments (Figure 3C and Supplementary Figure 2). These data thus confirmed that the presence of cationic polymers does not interfere with immunodetection assays and suggest that their use could significantly enhance detection of several EV-contained markers after immunocapture.

**Figure 3.**
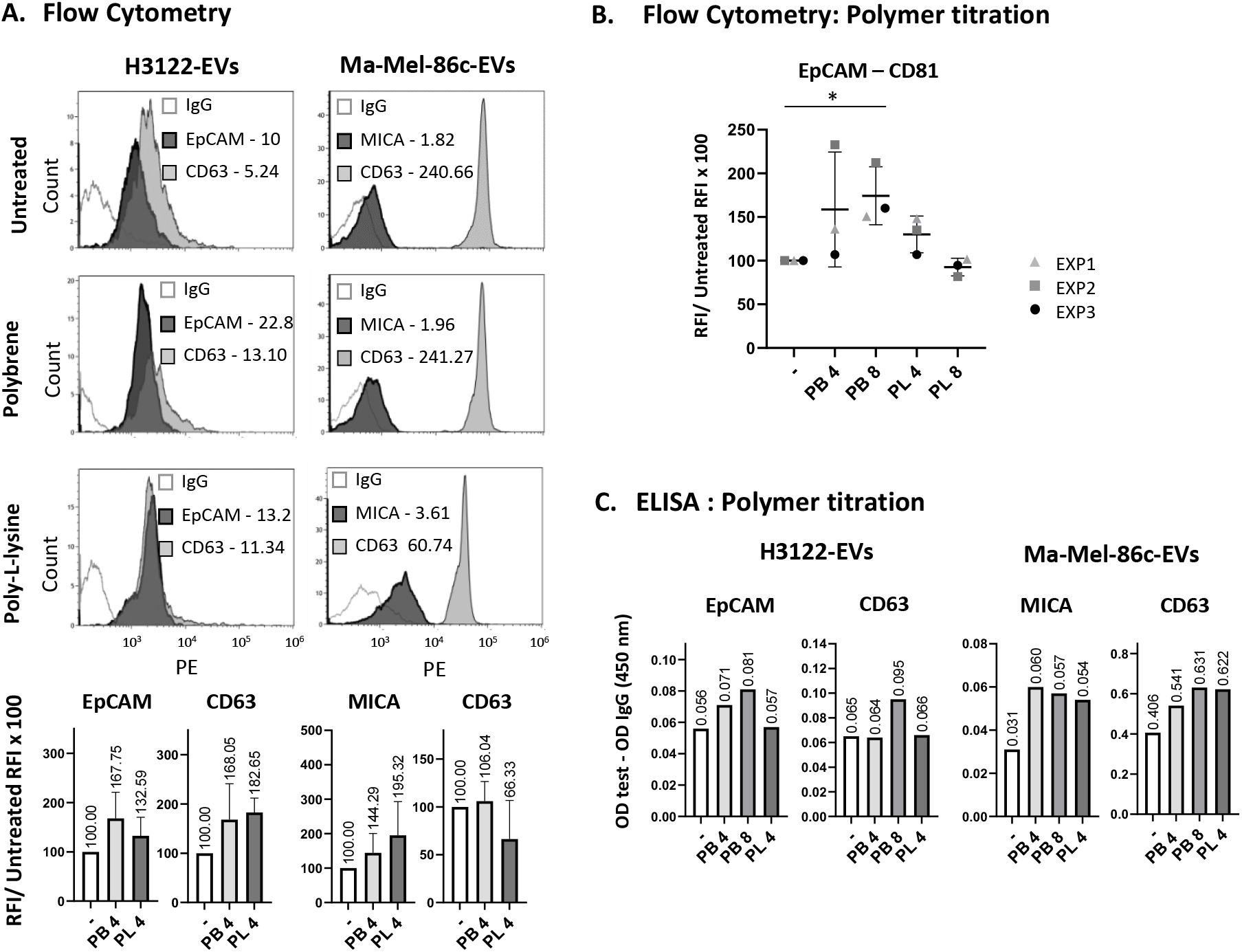
Cationic polymer addition increased EV detection by flow cytometry and ELISA. Lung cancer-derived EVs (from H3122 cell line) and melanoma-derived EVs (from Ma-Mel-86c cell line) were enriched by ultracentrifugation. 1 × 10^6^ H3122 EVs/µl or 1.8 × 10^7^ Ma-Mel-86c EVs/µl in 100 µl of PBS-Casein 1% were treated with 4 or 8 μg/ml polybrene (PB), 4 or 8 μg/ml poly-L-lysine (PL) or kept untreated for 18 h, as indicated in each panel. **A. Flow Cytometry**. EVs were incubated with 3000 of either anti-EpCAM, anti-MICA, anti-CD63 or IgG isotype control beads. Captured vesicles were detected by flow cytometry after incubation with anti-CD81-PE. Histograms from one representative experiment out of 3 are shown in the top panels. Relative Fluorescence Intensity 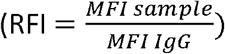 values are shown in each plot. Bar plots in the bottom panels represent the change in the RFI relative to the untreated condition. Mean and Standard Deviation from 3 independent experiments are represented. MFI: Median Fluorescence Intensity. **B. Titration of cationic polymers**. Tissue culture supernatant-derived EVs from the lung cancer cell line H3122 were incubated with either 3000 anti-EpCAM or IgG isotype control coated beads in 100 μl final volume of PBS-Casein 1%. EpCAM-captured vesicles were detected by flow cytometry after incubation with anti-CD81-PE. Relative increase of the RFI for EpCAM-CD81 detection, obtained in three experiment replicates (EXP), is shown. Statistical analysis was performed by a Two-way ANOVA Fisher’s LSD test. (* p < 0.05). **C. ELISA**. EVs enriched from tissue culture supernatant of the indicated cell lines were incubated in anti-EpCAM, anti-MICA or anti-CD63 antibody-coated plates. IgG Isotype-coated wells were used as negative control. Detection was performed with biotinylated anti-CD9 antibody followed by SA-HRP. Optical Density (OD) of the samples was represented after substraction of isotype OD. A representative experiment out of 3 is shown. Statistics from 3 independent experiments are available in Supplementary Figure 2.

### Optimised immunocapture of EVs allows detection of tetraspanins and tumour-associated proteins directly in plasma from cancer patients

The immunocapture experiments shown above strongly suggest that disruption of EV stability in suspension improves immunodetection by increasing EV availability to bind to antibody-coated surfaces. An alternative way to facilitate the encounter of microbeads and nanometric EVs during the capture step, would be to maximize the contact area by incubating the beads and EVs in a relatively broad surface with minimal volume. Thus, instead of using round-bottom tubes, in which beads accumulated in a relatively small diameter, flat-bottom micro-well plates were used (Supplementary Figure 3A). Indeed, reducing binding volumes extraordinarily increased detection of tetraspanins (Figure 4A, Supplementary Figure 3B). The great improvement in signal obtained in such small volumes, suggested that under these conditions this method might have enough sensitivity to directly detect EVs from biological samples. To test this hypothesis, antibodies against tetraspanins were used both for capture and detection in flow cytometry analysis of EVs directly in healthy donor plasma. CD63-CD81 EVs were efficiently captured and detected in plasma without any prior EV enrichment procedure, using as little as 12 µl of sample. Since, tumour-derived antigens are usually less abundant than tetraspanins on EVs, we also checked whether tumour-derived antigens in EVs could be directly detected in plasma using this methodology. As expected, healthy donor plasma was negative for EpCAM (Figure 4B, left). However, when H3122 lung cancer-derived EVs (positive for EpCAM) were added to a healthy donor plasma sample before the immunocapture experiment (at a final concentration of 6.25 × 10^7^ EVs/µl), the same sample showed a high positive signal for anti-EpCAM beads, as well as an increased signal for tetraspanins (Figure 4B, right). In order to estimate the amount of EVs that could be recognised using this method, a titration experiment was performed. The sensitivity of the assay was high enough to detect tumour associated antigens, such as EpCAM from H3122 lung cancer-derived EVs, at a concentration of 3.125 × 10^6^ EVs/μl in plasma (Supplementary Figure 4A).

**Figure 4:**
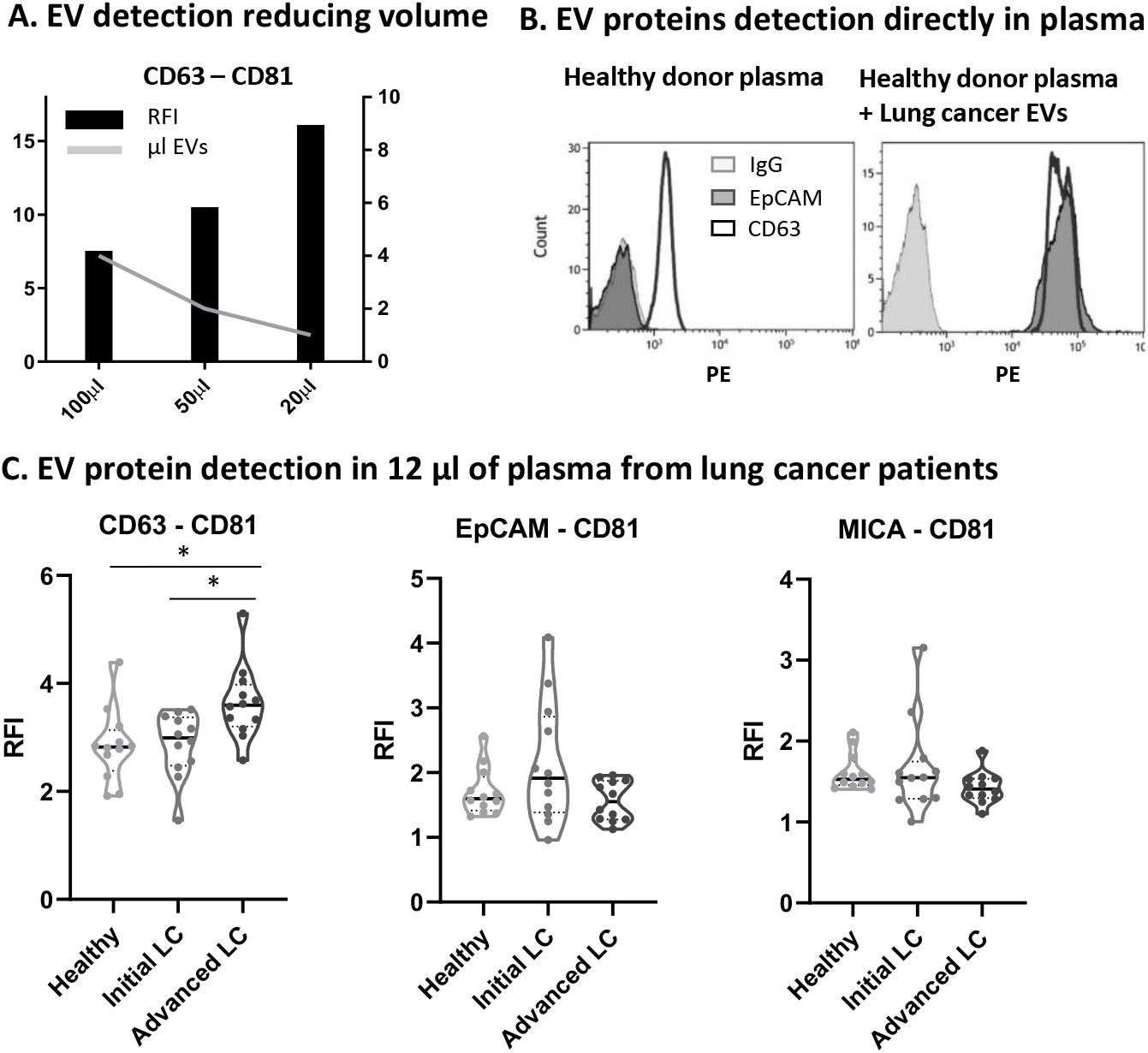
EV detection in reduced volume of plasma. A. Comparison of EV detection in decreasing volume. **A**. 3000 anti-CD63 beads were used per test in different incubation volumes (100 µL, 50 µl and 20 µl). Lung cancer-derived EVs (from H3122 cell line) were obtained by ultracentrifugation, incubated with beads for 16h and, after staining with anti-CD81-PE, washed and analysed by flow cytometry. The volume of EVs (5.35 ×10^8^ H3122 EVs/µl) added to each assay, is represented by a grey line (right Y-axe scale). Relative Fluorescence Intensity (RFI) obtained in each condition is represented in bars (left Y-axe scale). A representative experiment is shown out of 3 (available in Supplementary Figure 3). **B. Detection of EV proteins directly in plasma**. 12 µL of healthy donor plasma were incubated with 3000 anti-CD63, anti-EpCAM or IgG isotype control beads in a final volume of 25 µl. Captured EVs were detected with anti-CD81-PE. **Left**. Detection of EVs directly in 12 µl of healthy donor plasma. **Right**. 3.14 ×10^8^ H3122-derived EVs (from the H3122 cell line) were added to 12 µl of the same healthy donor plasma (2.6 × 10^7^ H3122 EVs/µl plasma). **C. EpCAM and MICA, can be detected directly in minimal volumes of plasma from a cohort of lung cancer (LC) patients by flow. cytometry** 12 µl of containing 3000 beads conjugated either with anti-CD63, anti-EpCAM or anti-MICA were incubated for 16 h with 12 µl of plasma (EDTA-tubes) from a cohort of lung cancer patients (12 initial stage and 12 advanced stage) and 12 healthy donors. The final volume of the assay was 24 µl. EVs captured in each assay were detected with anti-CD81-PE, or IgG-PE as a negative control to calculate the Relative Fluorescence Intensity (RFI). The mean RFI from three independent repetitions was calculated for each patient and represented as a violin plot for each group of patients. Statistical analysis was performed by one-way ANOVA (Tukey test for correction of multiple comparisons) or Krustal-Wallis non-parametric test (Dunns test for correction of multiple comparisons) with the same results. * p < 0.05, ** p < 0.01, *** p < 0.001, **** p < 0.0001). Patient samples were obtained at the University Hospital Clínica Universidad de Navarra.

The expression of candidate tumour-associated EV markers was then analysed in a pilot experiment using plasma from a small cohort of lung cancer patients compared to healthy donors. We used two tumour markers validated in the cell line-derived EVs: EpCAM, which has been described in EVs from epithelial cells and may be an epithelial cancer marker in blood (e.g. in Circulating Tumour Cells), and MICA, associated with cancer progression in many types of solid and haematological tumours ^38^. Plasma obtained from 12 lung cancer patients in initial stages, 12 patients with advanced lung cancer and 12 healthy donors was analysed (demographic and clinical data are available in Supplementary Table 1). The EVs present in 12 µl of plasma were captured either on anti-MICA, anti-EpCAM or anti-CD63-conjugated beads. Detection was performed with anti-CD81-PE or isotype-PE, as the negative control to calculate the RFI. CD63-CD81 positive EVs were efficiently detected in all samples (Figure 4C), with generally higher levels in cancer patients compared with healthy donors, being the difference only statistically significant for advanced stage lung cancer patients. EpCAM-CD81 positive EVs were higher in four initial stage lung cancer patients compared to healthy donors while MICA-CD81 positive EVs were observed to be higher in two initial stage lung cancer patients compared to healthy donors. Western Blot sensitivity did not allow detection of EpCAM in EV enriched preparations from the same donors (not shown) and only a faint band of MICA was visible for one of the two RFI positive patients in overexposed membranes (not shown).

### EV immunocapture combining small volume and cationic polymers in biological fluids has high sensitivity

We then analysed whether combining flat surfaces and minimal incubation volumes with cationic polymer addition further increased the signal detected in immunocapture assays using plasma from cancer patients. We first selected the best anti-coagulant for these assays (Supplementary Figure 5A). EDTA resulted in general in better signal, especially when polybrene was added. Comparison of the absence or presence of 8 µg/ml polybrene in these assays with minimal volumes of cancer patients’ plasma, confirmed a better signal when the cationic polymer was used (Supplementary Figure 5B). However, since in small volumes the interaction of the EVs with the beads is already enhanced, the increment obtained with polybrene was not as high as that observed in previous experiments performed in 100 μl (Supplementary Figure 4B) (148.5% vs 174.4% of the untreated signal), being only significant when the Confidence Interval (CI) was reduced from 95% to 90%.

Because our pilot experiment using plasma from lung cancer patients did not allow detection of tumour markers by WB, to estimate the sensitivity of this technique we used plasma from patients with other epithelial tumours where high expression of EpCAM in EVs has been reported ^39, 40^. Thus, to validate the methodology, this protein was studied together with tetraspanins, in plasma from 3 ovarian (Ov1-3) and 5 breast (Br1-5) cancer patients. 12 µl of plasma from each patient were captured on anti-EpCAM or anti-CD63-conjugated beads. Detection was performed with anti-CD81-PE, anti-CD9-PE or isotype-PE, as the negative control to calculate the RFI (Figure 5A). CD63-CD81 positive EVs were successfully detected in every sample, with higher signal in Ov1, Br1 and Br5. One ovarian cancer patient (Ov1) plasma was clearly positive for EpCAM^+^-CD81^+^ EVs, followed by a moderate signal in samples from Br1 and Br2 patients and lower signals were observed for the rest of the patients. Interestingly, at least in these sample types, the use of CD9 as a detection antibody was associated with higher RFIs than when EVs were detected using a CD81-specific mAb. Next, 200 µl of the same plasma samples were ultracentrifuged and the EV-enriched preparation was analysed in parallel by: 1) Western Blot (1/3 of EV prep volume) and 2) immunocapture followed by flow cytometry (1/10 of EV prep volume). The intensity of the CD81 bands detected by WB corresponded to the fluorescence intensities obtained by flow cytometry (in Figure 5B, compare WB bands with purple RFI heatmap). However, while flow cytometry detected different amounts of EpCAM in different patients (Figure 5B, turquoise blue RFI heatmap), only a long exposure of the membrane allowed a clear visualization of the EpCAM band in Ov1. Thus, flow cytometry detection of proteins in EVs is far more sensitive than WB.

**Figure 5:**
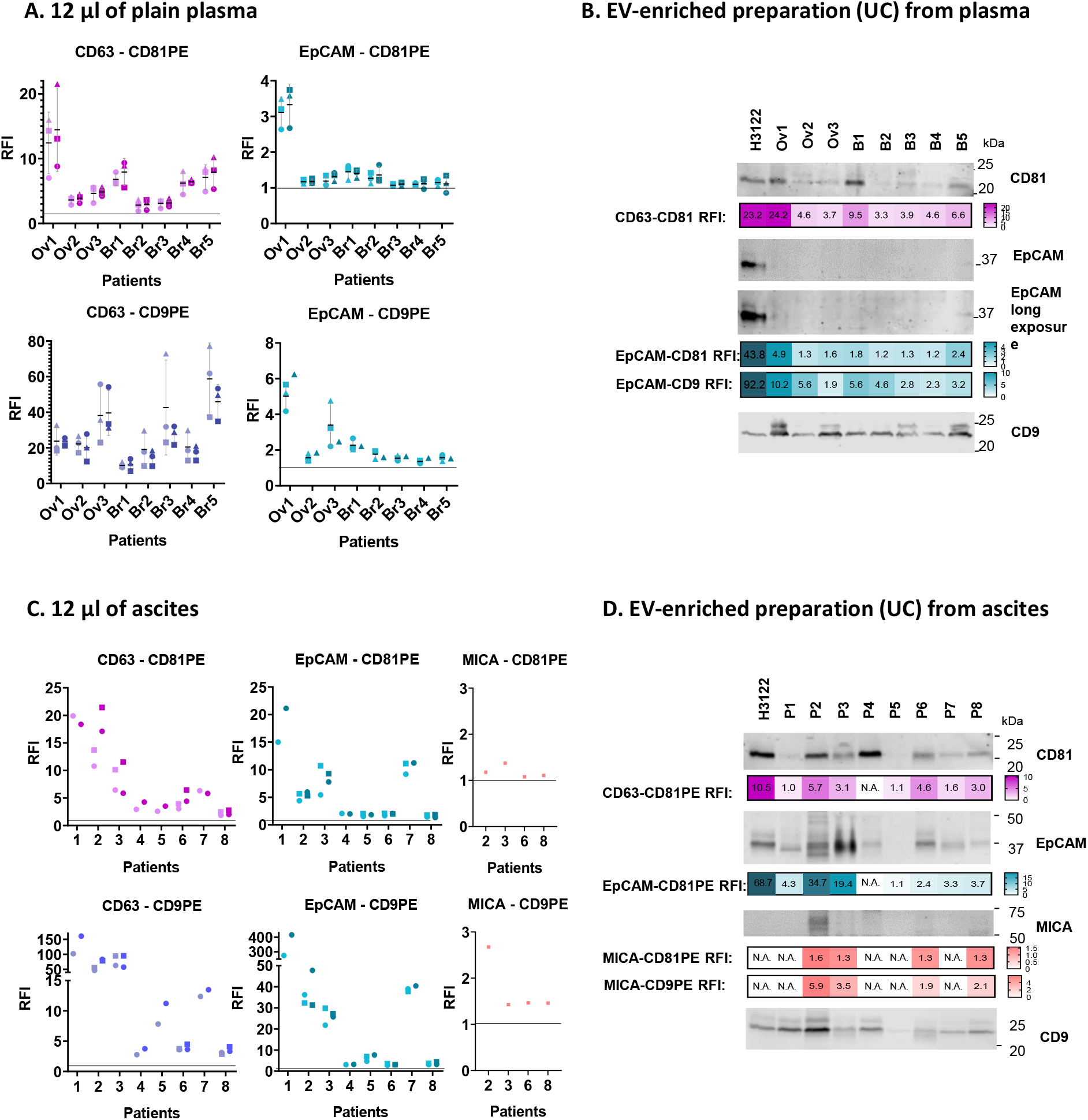
Tetraspanins and EV tumour-related proteins, EpCAM and MICA, can be detected directly in minimal volumes of plasma and ascitic fluid from cancer patients by flow cytometry, with better sensitivity than Western Blot. **Plasma (A, B). A**. 12 µl of PBS 1% casein containing 3000 beads conjugated with anti-CD63, anti-EpCAM or anti-MICA were incubated for 16 h with 12 µl of plasma (EDTA-tubes) from cancer patients (ovarian cancer Ov1-Ov3 and breast cancer Br1-Br5). Samples with addition of 8 µg/ml of polybrene were analysed in parallel (dark symbols). The final volume of the assay was 24 µl. EVs captured in each assay were detected with anti-CD81-PE, anti-CD9-PE or IgG-PE as a negative control to calculate the Relative Fluorescence Intensity (RFI). Mean and Standard Deviation of the RFI from 3 independent experiments (circles, squares and triangles) are represented. **B**. After ultracentrifugation of 200 µl of each plasma, the EV enriched preparation was resuspended in 15 µl. 5 µl of this EV preparation were loaded in a 12% SDS-PAGE gel and transferred to nitrocellulose. The membrane was immunoblotted for detection of EpCAM and MICA and the tetraspanins CD9 and CD81, as general EV markers. In parallel, 1.5 µl of the same EV preparation were incubated for 16 hours with 24 µl of PBS 1% casein containing 3000 beads conjugated with anti-CD63, anti-EpCAM or anti-MICA. EVs captured in each assay were detected by incubation with anti-CD81-PE or anti-CD9-PE. RFI values are represented as heatmaps (the number obtained is superimposed). EVs from the H3122 cell line were used as a control for EpCAM positive EVs (5.86 × 10^8^ particles/well for WB and 1.76 × 10^8^ particles/test for Flow Cytometry). **Ascites (C**,**D)**. Ovarian cancer patients ascites (1-8) were subjected to the same protocols as in A,B. Patient samples were obtained at the University Hospital Clínica Universidad de Navarra

Ascitic fluid from ovarian cancer patients has previously been reported to contain large amounts of EVs containing EpCAM ^41, 42^, and when 12 µl of ascites were tested by flow cytometry, EpCAM was successfully detected in all the ascitic fluid samples with high expression in several of them (Figure 5C). The tumour antigen MICA was also tested, and the EVs of one patient (P2) contained high amounts of this protein. Comparison with WB of EV enriched preparations confirmed the semi-quantitative power of the immunocapture technique (in figure 5D, compare the intensities of the bands with the RFI heatmaps). In ascites, it was possible to visualize EpCAM clearly by WB and the intensity of the observed band for each patient could be related with the fluorescence intensity observed in the flow cytometry heatmap (figure 5D, turquoise blue heatmap). Similarly, the detection of MICA in P2 by immunocapture correlated with the visualisation of a clear MICA band after WB analysis.

It is interesting to note that unprocessed ascites had somewhat different relative amounts of each protein marker when compared with the ultracentrifuged sample (eg. Patient 1 and 7 had high amounts of EpCAM in unprocessed plasma, but relatively lower in EVs after ultracentrifugation), highlighting the selection of different vesicle subpopulations during sequential centrifugation procedures.

In summary, direct immunocapture on 12 µl of patient plasma followed by flow cytometry yielded very sensitive results, compared to WB, and detection can be enhanced by the addition of cationic polymers. This procedure has the advantage of simple sample processing and eliminates the problems derived from sample manipulation, such as selection of EV subpopulations occurring after ultracentrifugation. The described EV characterization method may be applied to any biological fluid, such as urine and saliva as well as conditioned tissue culture supernatant (Supplementary Figure 6).

In conclusion, these experiments demonstrate that the optimised methodology for EV immunocapture can be used for a semi-quantitative analysis of EV tetraspanins as well as other proteins such as tumour-derived antigens, directly in 12 µl of plasma from cancer patients with minimal sample manipulation, opening the possibility for large screenings of multiple markers in patient cohorts.

## DISCUSSION

Immunocapture of EVs seems to yield, in general, lower detection levels than expected ^18, 43-45^. Here, we demonstrate that EVs behave as stable colloids with limited particle sedimentation, so that treatments predicted to modulate the biophysics of the colloidal suspension increase EV contact with functionalised surfaces and markedly improve EV protein detection after immunocapture. Based on this observation, a straight forward method for EV immunocapture was optimised for bead-assisted flow cytometry. Modifying the geometry of the reaction conditions, decreasing the incubation volume in a broad surface, also significantly increased assay sensitivity, so that conclusive results were obtained in plain plasma without the need to enrich EV samples. Detection could be further improved by the addition of cationic polymers. Thus, only 12 µl of plasma were enough for a clear detection of tetraspanins and other less abundant tumour markers by flow cytometry, with a sensitivity that is much higher than that of WB.

Although multiple previous reports have identified candidate disease markers in circulating EVs, their use in liquid biopsy still needs more definitive data. One of the biggest problems impeding marker validation is the difficulty of carrying out validation studies on large patient cohorts due to the limitation imposed by current EV enrichment methodology. By eliminating long manipulation protocols and increasing assay sensitivity, the method presented here could be used in high-throughput screenings in order to validate and discover new EV associated biomarkers. Further, these assays could be automatised in micro-titer plates, allowing standardization of the protocol.

The molecular basis of this enhanced immunocapture and detection can be explained by considering the physico-chemical characteristics of EVs as stable colloids. We hypothesized that EVs remain in suspension because gravity and buoyancy forces are not sufficient to counteract Brownian motion and electrostatic repulsion so that, in a bead-assisted assay, nanovesicles would remain in suspension while 6 µm beads precipitate relatively quickly, limiting the interaction of these particles. To test this hypothesis, polymer-induced colloidal flocculation was used. When cationic polymers were added to EV solutions, clusters of particles of different sizes were generated leading to higher sedimentation rates. These precipitation events correlated with an increase in protein detection. The cationic polymer-induced flocculation events occurring in this system could be due to particle aggregation after either neutralization of charges caused by adsorbed polyelectrolytes and/or formation of bridges between particles by simultaneous adsorption of polyelectrolyte chains onto more than two particles ^46-48^. Adding flocculants, allowed EVs to precipitate in a controlled manner leading to enhanced interaction with antibody-coated surfaces. Since charged polymers were used, the interaction is reversible and aggregates can be dissociated by successive washes, performed after the immunocapture step. This is a clear advantage compared to other polymers commonly used for EV precipitation, such as polyethylene glycol (PEG), a non-ionic uncharged hydrophilic polymer, whose water excluding properties create a high osmotic pressure causing irreversible protein precipitation in complex solutions ^49, 50^. In fact, DLS measurement of the mean diameter of an EV solution with 8% PEG 6000 was not possible, due to the presence of extensive polydispersity index (not shown). In contrast, due to the presence of charged groups, polyelectrolytes provide stronger and more tunable interactions and they are also sensitive to the solution pH and amount of electrolytes ^51^. In conclusion, these observations open a new avenue for further research on the interaction of polyelectrolytes with EV suspensions and biological fluids. Our data is also relevant for other applications involving EVs, such as isolation and recovery, in vitro EV-cell interaction, etc. ^52, 53^ and are in agreement with data from virus-containing solutions, where high molecular weight poly-L-lysine (up to 300 KDa) caused relatively higher aggregation than the lower molecular mass polymer polybrene (4-6 kDa) ^28^.

The findings with polymers motivated the adaptation of bead-assisted flow cytometry to enhance EV interaction with antibody-coated surfaces, and the use of minimal incubation volumes with optimised geometry, yielded markedly improved immunocapture results. The method was firstly tried with tumour-derived EV-enriched preparations from tissue culture supernatant and limits of detection were obtained in spiking experiments using plasma from healthy donors. Since our goal was to compare flow cytometry with other commonly used techniques, we focused on two putative cancer biomarkers that have been well characterised in our *in vitro* models. So, tetraspanins were analysed in plasma together with EpCAM, an epithelial marker commonly used to identify epithelial cancer circulating tumour cells. The immune activating molecule MICA belongs to a family of proteins, which are overexpressed on stressed cells such as tumour transformed or viral infected cells, and bind to the activating immune receptor NKG2D present on T lymphocytes and Natural Killer cells ^54, 55^. NKG2D-ligands have been shown to be overexpressed in most cancer cell lines and higher amounts of these soluble ligands in serum have been associated to worse cancer prognosis ^56-61^. MICA was studied because this molecule can be released from the cell surface in EVs ^62 63^.

In pilot experiments, since tumour staging and drug response can affect protein content in EVs, both initial and advanced lung cancer patient plasma were analysed. All samples were positive for CD63-CD81-EVs. Interestingly, advanced lung cancer patients had statistically significant higher levels of CD63-CD81-positive EVs compared with healthy donors and with initial stage patients. With respect to the detection by flow cytometry of tumour markers, EpCAM-CD81-containing EVs were efficiently detected in four initial stage lung cancer patients at higher levels than in healthy donor samples. MICA-CD81-containing EVs were also efficiently detected in two of these patients. According to published data, in some cases lung cancer patients EVs had low levels of EpCAM^39^. These data demonstrate the feasibility of our approach, but, of course, a large cohort needs to be studied for determination of cut-off values and for confirmation of the suitability of this particular tumour marker in diagnostics. Nevertheless, the finding of EpCAM-CD81-EVs in four initial stage patients, and not in advanced stage patients or healthy donors, agrees with previous observations that advanced tumour cells lose EpCAM expression as they undergo EMT (Epithelial to Mesenchymal Transition) ^64, 65^.

As we could not compare flow cytometry data with WB analysis in plasma from lung cancer patients, since no bands could be obtained for our molecules of interest, EpCAM and MICA, samples with higher expected content of these proteins on EVs, ovarian and breast cancer-derived plasma and ascitic fluid, were also analysed. Analysis of EpCAM in these samples clearly demonstrated the sensitivity and semi-quantitative capacity of the immunocapture assays. Detection by flow cytometry was considerably more sensitive than WB, since data could be obtained using three-fold less EV preparation than the amount loaded in WB. Further, we demonstrated that the intensity of fluorescence obtained in immunocapture followed by flow cytometry follows the same pattern as the intensity of the bands visualised by WB. These experiments demonstrate detection of the same specific molecule with similar relative intensity. The results also show that addition of cationic polymers can improve the signal obtained in small volumes of sample, confirming that the method could be used for large screenings of patients. Interestingly, abundant markers can lead to signal saturation and, in this case, polymer addition does not increase detection.

The data presented here, also demonstrated detection of EpCAM-CD81-containing EVs in small volumes of other biological fluids such as ascites from ovarian cancer patients, in general, in higher concentration than in plasma samples, in line with previous observations in breast cancer ^42^.

Another interesting observation is that the results of analyses of EVs prepared by ultracentrifugation from patient samples differ somewhat from the data obtained when EVs from unmanipulated plasma are assayed. This finding strongly suggests the loss or enrichment of different vesicle subpopulations during sample preparation, emphasising the necessity to use methods that allow EV characterization directly in biological samples and so avoid possible biases in the results obtained. Further, the selection of markers can affect results due to the relative abundance of tetraspanins in plasma or tumour cells as well as in different EV subpopulations ^18, 21^. For example, platelet derived EVs are devoid of CD81 but could contain CD9 ^66^. Here, EV heterogeneity could be observed when capturing with either CD63 or EpCAM and comparing the signal obtained for CD9 or CD81.

The data presented here demonstrate that EVs behave as colloidal suspensions, so, in immunoassays, the interphase contact with functionalised surfaces should be maximised. Cationic polymers can be combined with immunocapture methods directly in the biological sample to increase detection, without previous sample manipulation for EV enrichment. The described method should be useful to detect any EV protein of interest, since beads can be conjugated to any capture antibody to enrich and detect any EV subpopulation.

Altogether, the improved methodology has proven its potential to be used in high-throughput screenings of large cohorts of patients using multi-well plates and is adaptable to any laboratory setting. This will facilitate the validation and discovery of new body fluid EV-associated biomarkers, whose use can rapidly and easily be implemented in clinical settings.

## Author contributions

CCS, YCM, ESH, ASC generated and analysed data; provided material support and supervised the study. MVG, MYM, conceived, designed and supervised the study; MP, AGH, AR, ASC, ESH selected patient samples; MVG, MYM, AR, RJA provided material support; CCS and MVG wrote the manuscript with revisions from all authors.

## Acknowledgements

CCS and ESH are registered PhD student at the Universidad Autónoma de Madrid.

The authors thank Paolo Bergese (Institute for Research and Biomedical Innovation of Palermo - IRIB) for helpful discussions; A. Paschen (University Hospital Essen, Germany) for melanoma cell lines; V. Horejsi (Inst. of Molecular Genetics Academy of Sciences of the Czech Republic, Prague) for anti-CD81 antibody; Hector Peinado (National Centre for Oncological Research, CNIO, Spain) for the use of Nanosight NS500; MP Morales (Material Science Institute IICM-CSIC, Spain) for the use of Zetasizer NanoZS; C.A. Botello and J.R. Luque Ortega (laboratory of analytical ultracentrifugation and macromolecular interactions (LUAIM) at Centro de Investigaciones Biológicas Margarita Salas (CIB-CSIC)) for the analytic centrifuge service. R. Manzanero-Cancela (Spanish National Centre for Biotechnology CNB-CSIC) for technical help with EV preparations. Estíbalez Alegre (Clínica Universidad de Navarra) for help with patients’ samples selection. H.T. Reyburn (Spanish National Centre for Biotechnology CNB-CSIC) for helpful discussions and critical review of the manuscript. The Flow Cytometry Service and the Electron Microscopy Service at the Spanish National Centre for Biotechnology (CNB-CSIC).

## Funding

This work was supported by the [Spanish Ministry of Science and Innovation (MCIU/AEI/FEDER, EU) under Grants RTI2018-093569-B-I00, PID2020-119627GB-I00 and RED2018-102411-T (Translational Network for clinical application of EV, Tentacles); Madrid Regional Government under Grant “IMMUNOTHERCAN” (S2017/BMD-3733-2) and IND2019/BMD-17258.

## Conflict of Interest Disclosure

CCS, YCM, MYM, MVG, RJ are inventors on the European patent EP1641.1562. RJA is CEO of Immunostep, S.L. ABM is employed by Immunostep S.L. The rest of the authors declare no potential conflict of interest.

## Ethics approbal and patient consent statement

Experiments were carried out following the ethical principles established in the 1964 Declaration of Helsinki. Patients (or their representatives) were informed about the study and gave a written informed consent. This study used samples from 2 hospitals in Spain, Clínica Universidad de Navarra and Hospital Universitario Puerta de Hierro. Samples from Hospital Universitario Puerta de Hierro were obtained through the development of the research projects “PI17/01977” and “PIE14/00064”. Both projects were approved by the Hospital Puerta de Hierro Ethics Committee (internal code 79-18 and PI144, respectively). Collection of cancer patient samples at the Clínica Universidad de Navarra was approved by the Clínica Universidad de Navarra Ethics Committee 111/2010 “Estudio traslacional prospectivo de determinación de factores predictivos de eficacia y toxicidad en pacientes con cáncer” and included in the Spanish National Biobank Register with the code C.0003132 (Registro Nacional de Biobancos). A small cohort including 24 lung cancer patients and 12 healthy donors was approved by the project 2021.145.

## Data availability statement

All data generated in this study are included in this publication.

## Supplementary Figures

**Supplementary Figure 1**. Characterization of cell lines-derived EVs. **A**. Size and concentration analysis by Nanoparticle tracking analysis (NTA). Average size and concentration listed in the table were obtained in a Nanosight equipment capturing 3 videos of 60 s per measurement, with camera level 12, threshold 10 and temperature of 25 °C. Software NTA 3.1 (Malvern) was used for the analysis. ⍰ : diameter. **B**. Transmission Electron Microscopy (TEM) visualization. 1 µL EVs were diluted 1:10 in HBS and floated on a carbon-coated 400-mesh 240 Formvar grid, then incubated with 2% uranyl acetate and analysed using a Jeol JEM 1011 electron microscope operating at 245 100 kV with a CCD camera Gatan Erlangshen ES1000W. Pictures were taken at the Electron Microscopy Facility of the CNB. Bar: 100 nm. A representative image is shown. **C**. Protein marker characterization by Western Blot. EVs and whole cell lysates (L) were loaded in 12% SDS-PAGE gels. Membranes were immunoblotted for detection of: tetraspanins CD9, CD63, CD81 as general EV markers; β-actin as loading control; EpCAM and MICA as cancer-related markers; and calreticulin (CALR) as an endoplasmic reticulum resident protein not present in the EV fraction. Two gels were loaded: one gel, under non-reducing conditions and the other under reducing conditions, for actin detection. One representative experiment out of 3 is shown.

**Supplementary Figure 2. Cationic polymer addition increased cell lines-derived EV detection by ELISA**. 100 µl containing 1 × 10^6^ H3122 EVs/µl or 1.8 × 10^7^ Ma-Mel-86c EVs/µl in PBS-Casein 1% were treated with 4 or 8 μg/ml polybrene (PB), 4 μg/ml poly-L-lysine (PL) or kept untreated and incubated for 18 h in anti-EpCAM, anti-MICA or anti-CD63 antibody-coated plates. IgG coated wells were used as isotype control. EV detection was performed after incubation with biotinylated anti-CD9 antibody followed by SA-HRP. Optical Density (OD) was measured at 450 nm. After subtraction of the negative control, the binding was represented relative to the untreated sample. Statistics: one-way ANOVA Fisher’s LSD (* p <0.05; 95% confidence interval).

**Supplementary Figure 3. Volume reduction increased cell lines-derived EV detection. A**. Schematic representation of the binding surface. Diameter dimensions of the cytometry tube and the well a 96-well plate are depited to scale. Beads are represented in black and EVs in red. **B**. EV detection in different volumes. Different final volumes (100 µL, 50 µl and 20 µl) and different test conditions (5-ml tubes, flat bottom 96-well plates) were tested. Lung cancer-derived EVs (from H3122 cell line) were obtained by ultracentrifugation, incubated with beads for 18 h, stained with anti-CD81-PE and analysed by flow cytometry. Relative Fluorescence Intensity (RFI) obtained in each experiment from four replicates (EXP 1-4) are shown in the graph.

**Supplementary Figure 4. A.** Limit of detection of tetraspanins and EpCAM in plasma EVs, after addition of lung cancer-derived EVs. 12 µl of plasma samples containing decreasing concentrations of lung cancer-derived EVs were incubated with 12 µl of PBS-casein containing 3000 anti-CD63, anti-EpCAM beads or IgG isotype coated beads. Captured EVs were detected with anti-CD81-PE. Bar plots represent RFI (Relative Fluorescence Intensity) values obtained. The limit of detection for EpCAM was below 3.125 × 10^6^ EVs/μl. **B**. Titration of polymers for EpCAM detection in plasma after addition of lung cells-derived EVs. 3000 anti-EpCAM or IgG isotype-coated beads were incubated for 18 h with 2.6 × 10^6^ H3122-derived EVs/µl in 12 μL of healthy donor’s plasma in a final volume of 30 µL/test. Five different concentrations (0.5, 1, 2, 4 and 8 μg/ml) of Polybrene (PB) were compared to a polymer-untreated sample. Captured vesicles were detected by flow cytometry after incubation with anti-CD81-PE. IgG was used as a negative control. Increase of EpCAM-CD81 RFI relative to the untreated condition in four experiment replicates (EXP) is shown. Statistical analysis was performed by a Two-way ANOVA Fisher’s LSD test. (Confidence Interval CI = 90%) (* p < 0.1). Healthy donor samples were obtained at the University Hospital Puerta de Hierro.

**Supplementary Figure 5. Direct EpCAM and MICA detection in cancer patient plasma can be improved by the combination of small volume and cationic treatment. A**. Direct EpCAM and MICA detection in cancer patient plasma. 12 µl of PBS 1% casein containing 3000 beads conjugated with anti-CD63, anti-EpCAM or anti-MICA, as indicated, were incubated for 16 h with 12 µl of either serum (obtained in EDTA-tubes or heparin tubes) or plasma from each patient (cancer P1-P4 and non-cancer NC patients). The final volume of the assay was 26.5 µl, and two conditions were tested: either untreated EVs (in PBS 1% casein) (upper row) or treated with Polybrene at 8 µg/mL (lower row). EVs captured in each assay were detected with anti-CD81-PE. The signal obtained from incubation of plasma with IgG isotype control-coated beads was used to calculate the Relative Fluorescence Intensity (RFI). Mean and Standard Deviation from three independent experiments are represented. Statistical analysis was performed by a multiple t-test correcting for multiple comparisons by the Holm Sidack method (p < 0.05). * p < 0.05, ** p < 0.01, *** p < 0.001, **** p < 0.0001). Patient samples were obtained at the University Hospital Clínica Universidad de Navarra. **B**. 12 µl of PBS-casein containing 3000 beads conjugated with anti-CD63, anti-EpCAM or anti-MICA, as indicated, were incubated for 16 h with 12 µl of EDTA-plasma from each patient (cancer P1-P4 and non-cancer NC patients) either treated with Polybrene at 8 µg/mL or untreated with polymer. The final volume of the assay was 26.5 µl. Captured EVs were detected with anti-CD81-PE. The signal obtained from incubation of plasma with IgG isotype control-coated beads was used to calculate the Relative Fluorescence Intensity (RFI). Mean and Standard Deviation from three independent experiments are represented. Statistical analysis was performed by a multiple t-test correcting for multiple comparisons by the Holm Sidack method (p < 0.05). * p < 0.05, ** p < 0.01, *** p < 0.001, **** p < 0.0001). Patient samples were obtained at the University Hospital Clínica Universidad de Navarra.

**Supplementary Figure 6. Tetraspanins and EpCAM can be detected directly in minimal volumes of different biological fluids by flow cytometry. Polybrene can enhance low signals**. PBS 1% casein containing 3000 beads conjugated either with anti-CD63, anti-MICA or anti-EpCAM (as indicated) were incubated for 16 h with the indicated volumes of conditioned medium (**A**), urine (**B**) or saliva from a healthy donor (**C**), either untreated (PBS 1% casein) or treated with polybrene at 8 µg/mL. The final volume of the assay was 100 µl in A, and 26.5 µl in B, C. EVs captured in each assay were detected with anti-CD81-PE or anti-CD9-PE as indicated. Isotype-PE was used as a negative control to calculate the RFI: Relative Fluorescence Intensity. Samples were centrifuged once 10 min at 200 *x g* before analysis, except saliva which was centrifuged twice. Urine samples were obtained at the University Hospital Clínica Universidad de Navarra (P1 and P2 are male patients that required a urine test, P3-6 are prostate cancer patients).

## Notes

### Competing Interest Statement

CCS, YCM, MYM, MVG, RJ are inventors on the European patent EP1641.1562. RJ is CEO of Immunostep, S.L. AB is employed by Immunostep, S.L.

## References

1. Yanez-Mo M, Siljander PR, Andreu Z, et al. Biological properties of extracellular vesicles and their physiological functions. J Extracell Vesicles. 2015;4: 27066.

2. Bobrie A, Colombo M, Krumeich S, Raposo G, Thery C. Diverse subpopulations of vesicles secreted by different intracellular mechanisms are present in exosome preparations obtained by differential ultracentrifugation. J Extracell Vesicles. 2012;1.

3. Verweij FJ, Balaj L, Boulanger CM, et al. The power of imaging to understand extracellular vesicle biology in vivo. Nat Methods. 2021.

4. Yuana Y, Sturk A, Nieuwland R. Extracellular vesicles in physiological and pathological conditions. Blood Rev. 2013;27: 31–39.

5. Shah R, Patel T, Freedman JE. Circulating Extracellular Vesicles in Human Disease. N Engl J Med. 2018;379: 2180–2181.

6. Simeone P, Bologna G, Lanuti P, et al. Extracellular Vesicles as Signaling Mediators and Disease Biomarkers across Biological Barriers. Int J Mol Sci. 2020;21.

7. Tatischeff I. Current Search through Liquid Biopsy of Effective Biomarkers for Early Cancer Diagnosis into the Rich Cargoes of Extracellular Vesicles. Int J Mol Sci. 2021;22.

8. Pietrowska M, Zebrowska A, Gawin M, et al. Proteomic profile of melanoma cell-derived small extracellular vesicles in patients’ plasma: a potential correlate of melanoma progression. J Extracell Vesicles. 2021;10: e12063.

9. Hoshino A, Kim HS, Bojmar L, et al. Extracellular Vesicle and Particle Biomarkers Define Multiple Human Cancers. Cell. 2020;182: 1044–1061 e1018.

10. Gardiner C, Di Vizio D, Sahoo S, et al. Techniques used for the isolation and characterization of extracellular vesicles: results of a worldwide survey. J Extracell Vesicles. 2016;5: 32945.

11. Liangsupree T, Multia E, Riekkola ML. Modern isolation and separation techniques for extracellular vesicles. J Chromatogr A. 2021;1636: 461773.

12. Theodoraki MN, Hong CS, Donnenberg VS, Donnenberg AD, Whiteside TL. Evaluation of Exosome Proteins by on-Bead Flow Cytometry. Cytometry A. 2021;99: 372–381.

13. Liang Y, Lehrich BM, Zheng S, Lu M. Emerging methods in biomarker identification for extracellular vesicle-based liquid biopsy. J Extracell Vesicles. 2021;10: e12090.

14. Ibsen SD, Wright J, Lewis JM, et al. Rapid Isolation and Detection of Exosomes and Associated Biomarkers from Plasma. ACS Nano. 2017;11: 6641–6651.

15. Jeong S, Park J, Pathania D, Castro CM, Weissleder R, Lee H. Integrated Magneto-Electrochemical Sensor for Exosome Analysis. ACS Nano. 2016;10: 1802–1809.

16. Im H, Lee K, Weissleder R, Lee H, Castro CM. Novel nanosensing technologies for exosome detection and profiling. Lab Chip. 2017;17: 2892–2898.

17. Witwer KW, Buzas EI, Bemis LT, et al. Standardization of sample collection, isolation and analysis methods in extracellular vesicle research. J Extracell Vesicles. 2013;2.

18. Kowal J, Arras G, Colombo M, et al. Proteomic comparison defines novel markers to characterize heterogeneous populations of extracellular vesicle subtypes. Proc Natl Acad Sci U S A. 2016;113: E968–977.

19. Mathieu M, Nevo N, Jouve M, et al. Specificities of exosome versus small ectosome secretion revealed by live intracellular tracking of CD63 and CD9. Nat Commun. 2021;12: 4389.

20. Willms E, Cabanas C, Mager I, Wood MJA, Vader P. Extracellular Vesicle Heterogeneity: Subpopulations, Isolation Techniques, and Diverse Functions in Cancer Progression. Front Immunol. 2018;9: 738.

21. Jeppesen DK, Fenix AM, Franklin JL, et al. Reassessment of Exosome Composition. Cell. 2019;177: 428–445 e418.

22. Campos-Silva C, Suárez H, Jara-Acevedo R, et al. High sensitivity detection of extracellular vesicles immune-captured from urine by conventional flow cytometry. Sci Rep. 2019;9: 2042.

23. Morrison ID, Ross S. Colloidal dispersions : suspensions, emulsions, and foams. New York : Wiley-Interscience, 2002.

24. Stepto RFT. Dispersity in polymer science (IUPAC Recommendation 2009). Polymer International. 2010;59: 23–24.

25. Shboul AA, Pierre F, Claverie JP. Functional Materials: For Energy, Sustainable Development and Biomedical Sciences. In: Mario L, Robert G, editors. 2. A primer on polymer colloids: structure, synthesis and colloidal stability:: De Gruyter, 2014:9–36.

26. Tabujew I, Peneva K. CHAPTER 1 Functionalization of Cationic Polymers for Drug Delivery Applications. Cationic Polymers in Regenerative Medicine: The Royal Society of Chemistry, 2015:1-29. 27.

27. Murthy VS, Cha JN, Stucky GD, Wong MS. Charge-driven flocculation of poly(L-lysine)-gold nanoparticle assemblies leading to hollow microspheres. J Am Chem Soc. 2004;126: 5292–5299.

28. Davis HE, Rosinski M, Morgan JR, Yarmush ML. Charged polymers modulate retrovirus transduction via membrane charge neutralization and virus aggregation. Biophys J. 2004;86: 1234–1242.

29. Ugurel S, Thirumaran RK, Bloethner S, et al. B-RAF and N-RAS mutations are preserved during short time in vitro propagation and differentially impact prognosis. PLoS One. 2007;2: e236.

30. Zhao F, Sucker A, Horn S, et al. Melanoma Lesions Independently Acquire T-cell Resistance during Metastatic Latency. Cancer Res. 2016;76: 4347–4358.

31. Lopez-Cobo S, Pieper N, Campos-Silva C, et al. Impaired NK cell recognition of vemurafenib-treated melanoma cells is overcome by simultaneous application of histone deacetylase inhibitors. Oncoimmunology. 2018;7: e1392426.

32. Ashiru O, Boutet P, Fernandez-Messina L, et al. Natural killer cell cytotoxicity is suppressed by exposure to the human NKG2D ligand MICA*008 that is shed by tumor cells in exosomes. Cancer Res. 2010;70: 481–489.

33. Campos-Silva C, Caceres-Martell Y, Lopez-Cobo S, et al. An Immunocapture-Based Assay for Detecting Multiple Antigens in Melanoma-Derived Extracellular Vesicles. Methods Mol Biol. 2021;2265: 323-344. 34.

34. Thery C, Witwer KW, Aikawa E, et al. Minimal information for studies of extracellular vesicles 2018 (MISEV2018): a position statement of the International Society for Extracellular Vesicles and update of the MISEV2014 guidelines. J Extracell Vesicles. 2018;7: 1535750.

35. Lopez-Cobo S, Campos-Silva C, Moyano A, et al. Immunoassays for scarce tumour-antigens in exosomes: detection of the human NKG2D-Ligand, MICA, in tetraspanin-containing nanovesicles from melanoma. J Nanobiotechnology. 2018;16: 47.

36. Montis C, Zendrini A, Valle F, et al. Size distribution of extracellular vesicles by optical correlation techniques. Colloids Surf B Biointerfaces. 2017;158: 331–338.

37. Oliveira-Rodriguez M, Lopez-Cobo S, Reyburn HT, et al. Development of a rapid lateral flow immunoassay test for detection of exosomes previously enriched from cell culture medium and body fluids. J Extracell Vesicles. 2016;5: 31803.

38. Sharma P, Diergaarde B, Ferrone S, Kirkwood JM, Whiteside TL. Melanoma cell-derived exosomes in plasma of melanoma patients suppress functions of immune effector cells. Sci Rep. 2020;10: 92.

39. Wei P, Wu F, Kang B, et al. Plasma extracellular vesicles detected by Single Molecule array technology as a liquid biopsy for colorectal cancer. J Extracell Vesicles. 2020;9: 1809765.

40. Zhao Z, Yang Y, Zeng Y, He M. A microfluidic ExoSearch chip for multiplexed exosome detection towards blood-based ovarian cancer diagnosis. Lab Chip. 2016;16: 489–496.

41. Runz S, Keller S, Rupp C, et al. Malignant ascites-derived exosomes of ovarian carcinoma patients contain CD24 and EpCAM. Gynecol Oncol. 2007;107: 563–571.

42. Rupp AK, Rupp C, Keller S, et al. Loss of EpCAM expression in breast cancer derived serum exosomes: role of proteolytic cleavage. Gynecol Oncol. 2011;122: 437–446.

43. Clayton A, Court J, Navabi H, et al. Analysis of antigen presenting cell derived exosomes, based on immuno-magnetic isolation and flow cytometry. J Immunol Methods. 2001;247: 163–174.

44. Morales-Kastresana A, Jones JC. Flow Cytometric Analysis of Extracellular Vesicles. Methods Mol Biol. 2017;1545: 215–225.

45. Wiklander OPB, Bostancioglu RB, Welsh JA, et al. Systematic Methodological Evaluation of a Multiplex Bead-Based Flow Cytometry Assay for Detection of Extracellular Vesicle Surface Signatures. Front Immunol. 2018;9: 1326.

46. Lim VHY, Y.; Doan, Y.T.H.; Adachi, Y. Inhibition of Cationic Polymer-Induced Colloid Flocculation by Polyacrylic Acid. Water. 2018;10: 1215.

47. Adachi Y, Kobayashi A, Kobayashi M. Structure of Colloidal Flocs in relation to the Dynamic Properties of Unstable Suspension. International Journal of Polymer Science. 2012;2012: 574878.

48. Zhou Y, Franks GV. Flocculation Mechanism Induced by Cationic Polymers Investigated by Light Scattering. Langmuir. 2006;22: 6775–6786.

49. Effio CL, Hubbuch J. Next generation vaccines and vectors: Designing downstream processes for recombinant protein-based virus-like particles. Biotechnol J. 2015;10: 715–727.

50. Koho T, Mantyla T, Laurinmaki P, et al. Purification of norovirus-like particles (VLPs) by ion exchange chromatography. J Virol Methods. 2012;181: 6–11.

51. Khan N, Brettmann B. Intermolecular Interactions in Polyelectrolyte and Surfactant Complexes in Solution. Polymers (Basel). 2018;11.

52. Stranford DM, Hung ME, Gargus ES, Shah RN, Leonard JN. A Systematic Evaluation of Factors Affecting Extracellular Vesicle Uptake by Breast Cancer Cells. Tissue Eng Part A. 2017;23: 1274–1282.

53. Kim J, Lee H, Park K, Shin S. Rapid and Efficient Isolation of Exosomes by Clustering and Scattering. J Clin Med. 2020;9.

54. Fernandez-Messina L, Reyburn HT, Vales-Gomez M. Human NKG2D-ligands: cell biology strategies to ensure immune recognition. Front Immunol. 2012;3: 299.

55. Campos-Silva C. NKG2D-ligands: Putting everything under the same umbrella can be misleading 2018;91: 489–500.

56. Holdenrieder S, Stieber P, Peterfi A, Nagel D, Steinle A, Salih HR. Soluble MICA in malignant diseases. Int J Cancer. 2006;118: 684–687.

57. Holdenrieder S, Stieber P, Peterfi A, Nagel D, Steinle A, Salih HR. Soluble MICB in malignant diseases: analysis of diagnostic significance and correlation with soluble MICA. Cancer Immunol Immunother. 2006;55: 1584–1589.

58. Zhao Y, Chen N, Yu Y, et al. Prognostic value of MICA/B in cancers: a systematic review and meta-analysis. Oncotarget. 2017;8: 96384–96395.

59. Li K, Mandai M, Hamanishi J, et al. Clinical significance of the NKG2D ligands, MICA/B and ULBP2 in ovarian cancer: high expression of ULBP2 is an indicator of poor prognosis. Cancer Immunol Immunother. 2009;58: 641–652.

60. Jinushi M, Takehara T, Tatsumi T, et al. Impairment of natural killer cell and dendritic cell functions by the soluble form of MHC class I-related chain A in advanced human hepatocellular carcinomas. J Hepatol. 2005;43: 1013–1020.

61. Liu G, Lu S, Wang X, et al. Perturbation of NK cell peripheral homeostasis accelerates prostate carcinoma metastasis. J Clin Invest. 2013;123: 4410–4422.

62. Ashiru O, Lopez-Cobo S, Fernandez-Messina L, et al. A GPI anchor explains the unique biological features of the common NKG2D-ligand allele MICA*008. Biochem J. 2013.

63. Lopez-Cobo S. Glycosyl-Phosphatidyl-Inositol (GPI)-Anchors and Metalloproteases: Their Roles in the Regulation of Exosome Composition and NKG2D-Mediated Immune Recognition 2016;4: 97.

64. Keller L, Werner S, Pantel K. Biology and clinical relevance of EpCAM. Cell Stress. 2019;3: 165-180.65.

65. Byers LA, Diao L, Wang J, et al. An epithelial-mesenchymal transition gene signature predicts resistance to EGFR and PI3K inhibitors and identifies Axl as a therapeutic target for overcoming EGFR inhibitor resistance. Clin Cancer Res. 2013;19: 279–290.

66. Koliha N, Wiencek Y, Heider U, et al. A novel multiplex bead-based platform highlights the diversity of extracellular vesicles. J Extracell Vesicles. 2016;5: 29975.

