## Supplemental Figures and table for "A simple immunoassay for extracellular vesicle liquid biopsy in microliters of non-processed plasma"

**A. NTA**

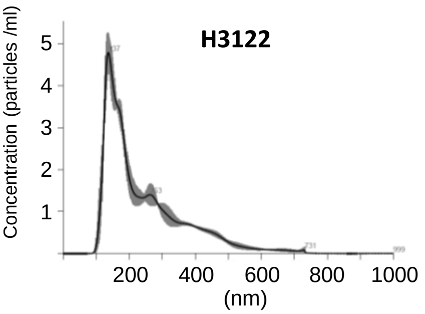


| | |
|---|---|
| Average ϕ | 244.2 +/- 6.1 nm |
| Mode ϕ | 139.2 +/- 3.1 nm |
| Particles/µL | $4.19 \cdot 10^8$ |

**B. TEM**

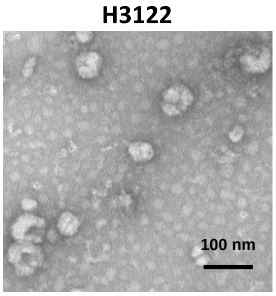


**C. WB**

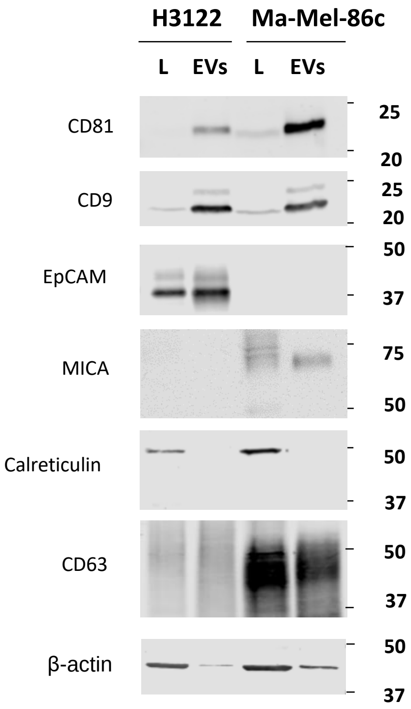


**Characterization of cell lines-derived EVs. A. Size and concentration analysis by Nanoparticle tracking analysis (NTA).** Average size and concentration listed in the table were obtained in a Nanosight equipment capturing 3 videos of 60 s per measurement, with camera level 12, threshold 10 and temperature of 25 °C. Software NTA 3.1 (Malvern) was used for the analysis. ϕ: diameter. **B. Transmission Electron Microscopy (TEM) visualization.** 1 µL EVs were diluted 1:10 in HBS and floated on a carbon-coated 400-mesh 240 Formvar grid, then incubated with 2% uranyl acetate and analysed using a Jeol JEM 1011 electron microscope operating at 245 100 kV with a CCD camera Gatan Erlangshen ES1000W. Pictures were taken at the Electron Microscopy Facility of the CNB. Bar: 100 nm. A representative image is shown. **C. Protein marker characterization by Western Blot.** EVs and whole cell lysates (L) were loaded in 12% SDS-PAGE gels. Membranes were immunoblotted for detection of: tetraspanins CD9, CD63, CD81 as general EV markers; β-actin as loading control; EpCAM and MICA as cancer-related markers; and calreticulin (CALR) as an endoplasmic reticulum resident protein not present in the EV fraction. Two gels were loaded: one gel, under non-reducing conditions and the other under reducing conditions, for actin detection. One representative experiment out of 3 is shown.

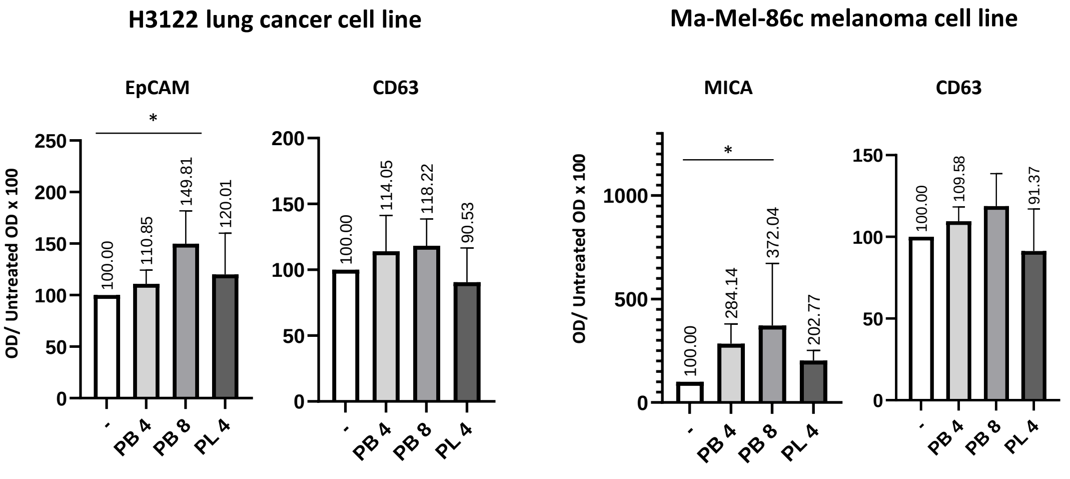


**Cationic polymer addition increased cell lines-derived EV detection by ELISA.** 100 µl containing 1 x $10^6$ H3122 EVs/µl or 1.8 x $10^7$ Ma-Mel-86c EVs/µl in PBS-Casein 1% were treated with 4 or 8 µg/ml polybrene (PB), 4 µg/ml poly-L-lysine (PL) or kept untreated and incubated for 18 h in anti-EpCAM, anti-MICA or anti-CD63 antibody-coated plates. IgG coated wells were used as isotype control. EV detection was performed after incubation with biotinylated anti-CD9 antibody followed by SA-HRP. Optical Density (OD) was measured at 450 nm. After subtraction of the negative control, the binding was represented relative to the untreated sample. Statistics: one-way ANOVA Fisher's LSD (* $p < 0.05$; 95% confidence interval).

**A. Schematic representation of the binding surface**

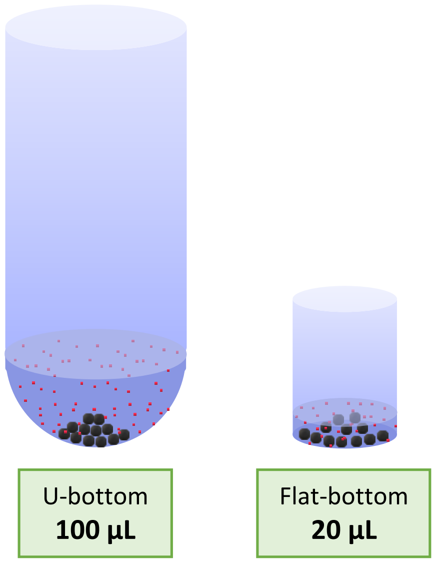


**B. EV detection using different test volume**

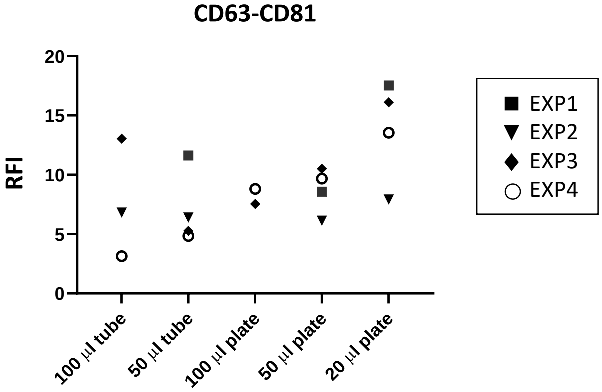


**Volume reduction increased cell lines-derived EV detection. A. Schematic representation of the binding surface.** Diameter dimensions of the cytometry tube and the well a 96-well plate are depited to scale. Beads are represented in black and EVs in red. **B. EV detection in different volumes.** Different final volumes (100 µL, 50 µl and 20 µl) and different test conditions (5-ml tubes, flat bottom 96-well plates) were tested. Lung cancer-derived EVs (from H3122 cell line) were obtained by ultracentrifugation, incubated with beads for 18 h, stained with anti-CD81-PE and analysed by flow cytometry. Relative Fluorescence Intensity (RFI) obtained in each experiment from four replicates (EXP 1-4) are shown in the graph.

**A. Limit of detection for EV tumour antigens in plasma**

**B. Polymers titration in plasma**

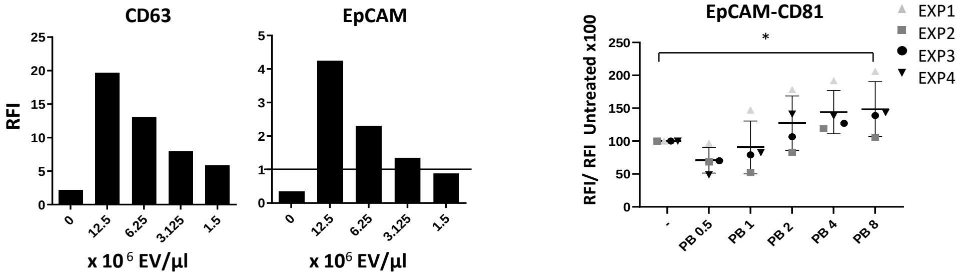


**A. Limit of detection of tetraspanins and EpCAM in plasma EVs, after addition of lung cancer-derived EVs.** 12 µl of plasma samples containing decreasing concentrations of lung cancer-derived EVs were incubated with 12 µl of PBS-casein containing 3000 anti-CD63, anti-EpCAM beads or IgG isotype coated beads. Captured EVs were detected with anti-CD81-PE. Bar plots represent RFI (Relative Fluorescence Intensity) values obtained. The limit of detection for EpCAM was below 3.125 x $10^6$ EVs/µl. **B. Titration of polymers for EpCAM detection in plasma after addition of lung cells-derived EVs .** 3000 anti-EpCAM or IgG isotype-coated beads were incubated for 18 h with 2.6 x $10^6$ H3122-derived EVs/µl in 12 µL of healthy donor's plasma in a final volume of 30 µL/test. Five different concentrations (0.5, 1, 2, 4 and 8 µg/ml) of Polybrene (PB) were compared to a polymer-untreated sample. Captured vesicles were detected by flow cytometry after incubation with anti-CD81-PE. IgG was used as a negative control. Increase of EpCAM-CD81 RFI relative to the untreated condition in four experiment replicates (EXP) is shown. Statistical analysis was performed by a Two- way ANOVA Fisher's LSD test. (Confidence Interval CI = 90%) (* $p < 0.1$). Healthy donor samples were obtained at the University Hospital Puerta de Hierro.

### A. Direct EV detection in plasma using different anti-coagulants

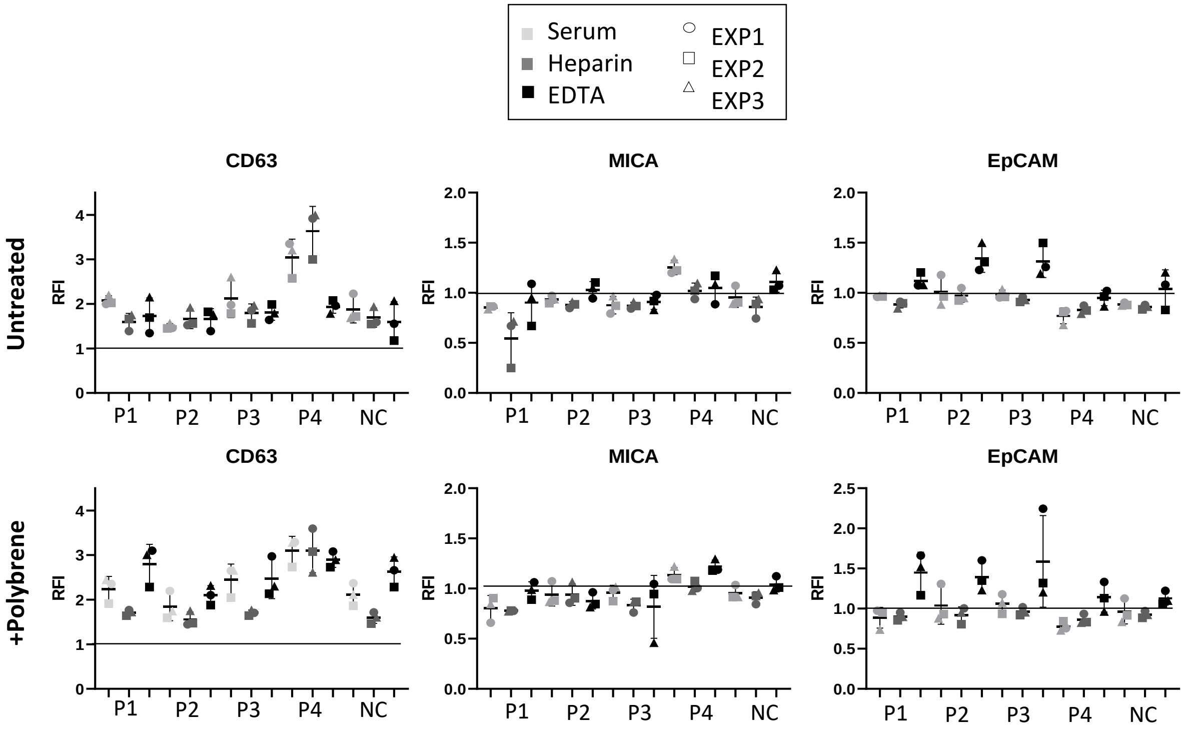


### B. Combination of small volume and cationic treatment

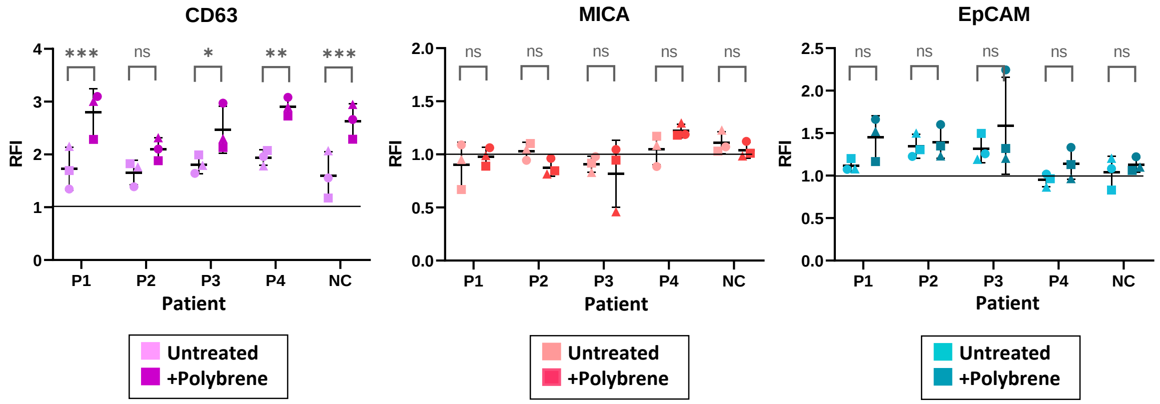


**Direct EpCAM and MICA detection in cancer patient plasma can be improved by the combination of small volume and cationic treatment**. **A. Direct EpCAM and MICA detection in cancer patient plasma.** 12 µl of PBS 1% casein containing 3000 beads conjugated with anti-CD63, anti-EpCAM or anti-MICA, as indicated, were incubated for 16 h with 12 µl of either serum (obtained in EDTA-tubes or heparin tubes) or plasma from each patient (cancer P1-P4 and non-cancer NC patients). The final volume of the assay was 26.5 µl, and two conditions were tested: either untreated EVs (in PBS 1% casein) (upper row) or treated with Polybrene at 8 µg/mL (lower row). EVs captured in each assay were detected with anti-CD81-PE. The signal obtained from incubation of plasma with IgG isotype control-coated beads was used to calculate the Relative Fluorescence Intensity (RFI). Mean and Standard Deviation from three independent experiments are represented. Statistical analysis was performed by a multiple t-test correcting for multiple comparisons by the Holm Sidack method ($p < 0.05$). * $p < 0.05$, ** $p < 0.01$, *** $p < 0.001$, **** $p < 0.0001$). Patient samples were obtained at the University Hospital Clínica Universidad de Navarra. **B.** 12 µl of PBS-casein containing 3000 beads conjugated with anti-CD63, anti-EpCAM or anti-MICA, as indicated, were incubated for 16 h with 12 µl of EDTA-plasma from each patient (cancer P1-P4 and non-cancer NC patients) either treated with Polybrene at 8 µg/mL or untreated with polymer. The final volume of the assay was 26.5 µl. Captured EVs were detected with anti-CD81-PE. The signal obtained from incubation of plasma with IgG isotype control-coated beads was used to calculate the Relative Fluorescence Intensity (RFI). Mean and Standard Deviation from three independent experiments are represented. Statistical analysis was performed by a multiple t-test correcting for multiple comparisons by the Holm Sidack method ($p < 0.05$). * $p < 0.05$, ** $p < 0.01$, *** $p < 0.001$, **** $p < 0.0001$). Patient samples were obtained at the University Hospital Clínica Universidad de Navarra.

### Supplementary Figure 6

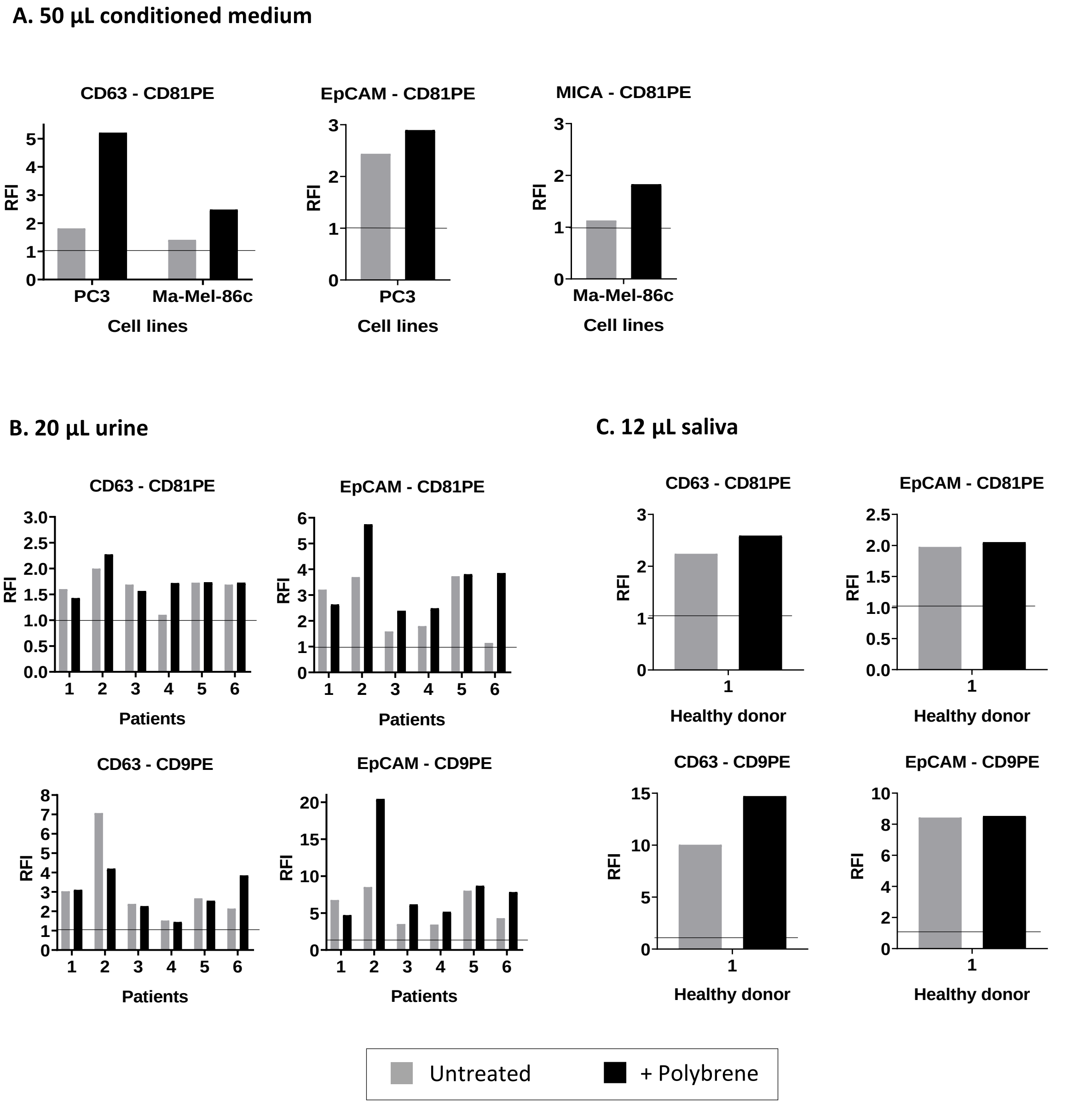


**Tetraspanins and EpCAM can be detected directly in minimal volumes of different biological fluids by flow cytometry. Polybrene can enhance low signals.** PBS 1% casein containing 3000 beads conjugated either with anti-CD63, anti-MICA or anti-EpCAM (as indicated) were incubated for 16 h with the indicated volumes of conditioned medium (A), urine (B) or saliva from a healthy donor (C), either untreated (PBS 1% casein) or treated with polybrene at 8 µg/mL. The final volume of the assay was 100 µl in A, and 26.5 µl in B, C. EVs captured in each assay were detected with anti-CD81-PE or anti-CD9-PE as indicated. Isotype-PE was used as a negative control to calculate the RFI: Relative Fluorescence Intensity. Samples were centrifuged once 10 min at 200 *x g* before analysis, except saliva which was centrifuged twice. Urine samples were obtained at the University Hospital Clínica Universidad de Navarra (P1 and P2 are male patients that required a urine test, P3-6 are prostate cancer patients).

**Supplementary Table 1. Lung cancer patients. Demographic and pathological data**

| Characteristics | | Initial stage lung cancer (N=12) | Advanced stage lung cancer (N=12) | Healthy donors (N=12) |
|---|---|---|---|---|
| | | Number (%) | Number (%) | Number (%) |
| *Age* | Mean | 68.83 | 66.08 | 38.66 |
| | Min-Max | 51-87 | 45-78 | 22-56 |
| *Sex* | Male | 8 (66) | 7 (58.33) | 3 (25) |
| | Female | 4 (33) | 5 (41.66) | 9 (75) |
| *Histology* | **Non-Small Cell Lung Cancer (NSCLC)** | 9 (75) | 9 (75) | |
| | - Adenocarcinoma | - 8 (66.66) | - 8 (66.66) | |
| | - Squamous cell carcinoma | - 1 (8.33) | - 1 (8.33) | |
| | - Large cell carcinoma | - 0 (0) | - 0 (0) | |
| | **Small cell lung cancer (SCLC)** | 1 (8.33) | 2 (16.66) | |
| | **Other (pleural mesothelioma or epidermoid carcinoma)** | 2 (16) | 1 (8.33) | |
| *Clinical Stage* | IA | 3 (25) | | |
| | IB | 2 (16.66) | | |
| | IIA | 1 (8.33) | | |
| | IIB | 6 (50) | | |
| | IIIA | | 2 (16.66) | |
| | IIIB | | 2 (16.66) | |
| | IV | | 8 (66.66) | |
